# The tumor suppressor CYLD acts as a deubiquitinase for mTOR to constrain its activity

**DOI:** 10.1101/2025.09.01.673523

**Authors:** Stephanie A. Fernandes, Jiyoung Pan, Diana S. Terziyska, Seda Koyuncu, Xiaolei Ding, István Balázs Németh, Sabine Wilhelm, Julian Nüchel, Suzan Al-Gburi, Christos Gonidas, Manolis Pasparakis, George Mosialos, Márta Széll, Aurelio A. Teleman, Sabine A. Eming, David Vilchez, Constantinos Demetriades

## Abstract

Proper control of mTOR (mechanistic/mammalian target of rapamycin) signaling is relevant for health, disease and ageing. Information from intra- and extra-cellular signaling cues is transmitted to mTOR through an intricate signaling network that impinges on the Rag and Rheb GTPases to regulate its localization and activity. Interestingly, although mTOR is a heavily ubiquitinated protein, the role of this post-translational modification (PTM) in regulating its activation status remains poorly understood. Here, through an unbiased RNAi screen, we identified the tumor suppressor CYLD deubiquitinase (DUB) as a direct negative regulator of both mTORC1 and mTORC2 activities. Mechanistically, CYLD interacts with mTOR and removes non-degradative, K63-linked ubiquitin (Ub) chains from multiple of its residues. Consequently, CYLD loss-of-function cells are characterized by mTORC1/2 hyperactivation, elevated rates of protein synthesis, increased cell size, and resistance to serum-starvation-induced activation of cell death pathways. Moreover, silencing of *cyld-1*, the *C. elegans* CYLD ortholog, fully reverses the extended lifespan of low- TORC1-activity mutant worms. Finally, we find that inactivation of CYLD is associated with hyperactivation of mTORC1 also in skin biopsies from *CYLD* cutaneous syndrome (CCS) patients. In sum, our findings highlight CYLD as a sentinel of mTOR hyperactivation via direct control of its ubiquitination, and suggest that dysregulated mTOR activity may contribute to the development and progression of CCS tumors.

## Introduction

Signaling pathways are essential for conveying and amplifying messages that are required for cells to dynamically adapt their function to the conditions in their environment ^1, 2^, while dysregulation of these molecular mechanisms can have life- threatening consequences ^3^. Signal transmission commonly occurs through the addition of post-translational modifications (PTMs) to specific proteins that participate in these signaling cascades. As such, PTMs act as dynamic molecular switches, influencing various properties of individual proteins and protein complexes in diverse physiological and pathophysiological contexts ^4–9^. A remarkable example is the mTOR signaling pathway, comprised of dozens of proteins that are modified by PTMs, like phosphorylation and ubiquitination, impacting their activity, localization and stability ^10,11^. Hence, the mechanisms involved in the addition and removal of PTMs from key signaling components are central to the understanding of cellular homeostasis in health, disease, and ageing.

The mTOR kinase is recognized as the master regulator of cellular physiology, serving as the core component of two multiprotein signaling complexes, mTORC1 (mTOR complex 1) and mTORC2. mTORC1 functions as a central regulatory hub, orchestrating the activation of key metabolic and other growth-related biosynthetic cellular processes, including protein synthesis ^12^. Conversely, mTORC2 plays a pivotal role in regulating cell proliferation, cytoskeletal organization, and cell survival ^13^. Notably, both mTOR complexes integrate signals from intra- and extracellular stimuli, like nutrient sufficiency and growth factor availability, translating these inputs into molecular and metabolic adaptations that enable cells to respond appropriately. The essential role of mTOR signaling in maintaining cellular homeostasis is evident in its dysregulation, which has been implicated in ageing, cancer, as well as in several metabolic and neurological disorders in humans ^12^.

mTORC1 localization and activity is under the control of a set of small GTPases, primarily the Rag and Rheb GTPases, acting downstream of a vast signaling network that relays information about the nutritional, hormonal and stress status of a cell ^12, 14, 15^, while the regulation of mTORC2 in response to growth factor stimulation involves its PI(3,4,5)P_3_ (phosphatidylinositol 3,4,5-triphosphate)-dependent recruitment to the plasma membrane ^13^. An additional layer of mTORC1/2 regulation involves the post-translational modification of mTOR itself, including phosphorylation, ubiquitination, and acetylation ^11, 16–18^. Notably, although mTOR is a heavily ubiquitinated protein, the precise role of these modifications on its function remains so far poorly understood. Cumulative evidence has shed light into the E3 Ub ligases that mediate mTOR ubiquitination: for instance, the tumor suppressor FBXW7 (F-box/WD repeat-containing protein 7) controls mTOR protein levels by facilitating its ubiquitination and subsequent degradation ^19^. Furthermore, amino acid (AA) stimulation promotes the activation of mTORC1 also via the addition of non-degradative, K63-linked Ub chains to mTOR, a process that depends on the p62/SQSTM1 (sequestosome 1) adaptor protein and catalyzed by the TRAF6 E3 Ub ligase ^20^. In contrast, AA insufficiency was previously reported to cause the FBXO22-mediated attachment of K27-linked Ub chains at the K2066 residue of mTOR ^21^. The same mTOR residue can be also ubiquitinated by a different enzyme, the PRKN (parkin RBR E3 ubiquitin protein ligase), in response to mitochondrial stress ^22^. Altogether, these previous studies underscore the essential involvement of ubiquitination by multiple E3 ligase enzymes in the regulation of mTOR function. However, as protein ubiquitination is a reversible modification, the removal of Ub chains catalyzed by the respective DUB enzymes must be of equal importance for the fine-tuning of mTOR ubiquitination and activity. Notably, a DUB enzyme that acts directly on mTOR to regulate its activation status is not known.

Many DUB enzymes recognize single Ub molecules or Ub chains independently of the target to which they are attached. In contrast, the USP (ubiquitin-specific protease) family of DUBs is distinctive in that many of its members have the capacity to also engage in protein-protein interactions with their substrates ^23^. In particular, CYLD (CYLD Lysine 63 Deubiquitinase; also referred to as cylindromatosis) is a remarkable example of a DUB that directly interacts with its targets ^24–27^ and shows strong specificity towards the recognition and removal of K63- and M1-linked Ub chains ^28^. Highlighting its important role in cell growth and survival, CYLD was identified as a tumor suppressor. More specifically, inactivating mutations in the *CYLD* gene locus are causally associated with *CYLD* cutaneous syndrome (CCS) (also referred to as cylindromatosis or Brooke-Spiegler syndrome, BSS), a rare autosomal dominant human condition characterized by multiple tumors in the head and neck area that develop from structures associated with skin appendages ^29^.

Here, we report that CYLD directly deubiquitinates mTOR to keep its activity in check. Therefore, in cellular models of CYLD deficiency, the activities of mTORC1/2 and downstream physiological functions that are controlled by these complexes are found dysregulated. In addition, we reveal a role of CYLD in *C. elegans* ageing via the regulation of mTOR activity, and find that mTORC1 signaling is aberrantly elevated in skin biopsies of CCS/cylindromatosis patients. Collectively, our findings establish a tight connection between the tumor suppressor CYLD DUB and the growth-related kinase mTOR, and highlight mTOR deubiquitination by CYLD as an important layer of its regulation with potential relevance for ageing and disease.

## Results

### An unbiased RNAi screen highlights CYLD as a negative regulator of mTORC1

To identify DUBs that may act as putative mTORC1 regulators, we set up an unbiased RNAi screen for approximately 100 DUB-encoding genes in human breast cancer MCF-7 cells, using S6K (ribosomal protein S6 kinase) phosphorylation as a read-out for mTORC1 activity (Fig. 1a and Table S1; see also *Methods*). As controls, we included siRNA reagents targeting known mTORC1 pathway components such as *TSC2* (tuberous sclerosis complex 2; an upstream negative regulator of mTORC1), *AKT1* (AKT serine/threonine kinase 1; an upstream positive regulator of mTORC1), *RPTOR* (RAPTOR, regulatory-associated protein of mTOR; a core mTORC1 component), and S6K itself (encoded by the *RPS6KB1* gene). Confirming the robustness of our approach, knockdown of *S6K1*, *AKT1*, or *RAPTOR* led to a stark decrease in mTORC1 activity score, while knockdown of *TSC2* caused strong mTORC1 hyperactivation, as expected (Fig. 1b and Table S1). Knockdown of several DUB genes affected mTORC1 activity, positively or negatively (Table S1). Among the top hits that acted as putative negative regulators of mTORC1, we identified the tumor suppressor *CYLD* (CYLD lysine 63 deubiquitinase) gene, whose knockdown caused a similar upregulation of mTORC1 activity to that of *TSC2* knockdown (Fig. 1b and Table S1). Notably, inactivating germline mutations in *CYLD* are associated with CCS/cylindromatosis, a rare condition characterized by the growth of multiple benign tumors of the skin appendages in humans ^29, 30^. Given the well-established role of mTOR in supporting cancer cell growth ^31–33^, we hypothesized that inactivation of CYLD may be driving tumor growth, at least in part, via the hyperactivation of mTORC1 in cells. In support of this hypothesis, mutation analysis in diverse human cancers using the cBioPortal platform for cancer genomics (www.cbioportal.org) showed that inactivating mutations or genetic deletions in the *CYLD* locus do not coincide with such alterations in the *TSC2*, *TSC1*, or *FLCN* (folliculin) genes, suggesting that CYLD may be also acting as a negative upstream regulator of mTORC1 (Fig. S1).

**Figure 1.**
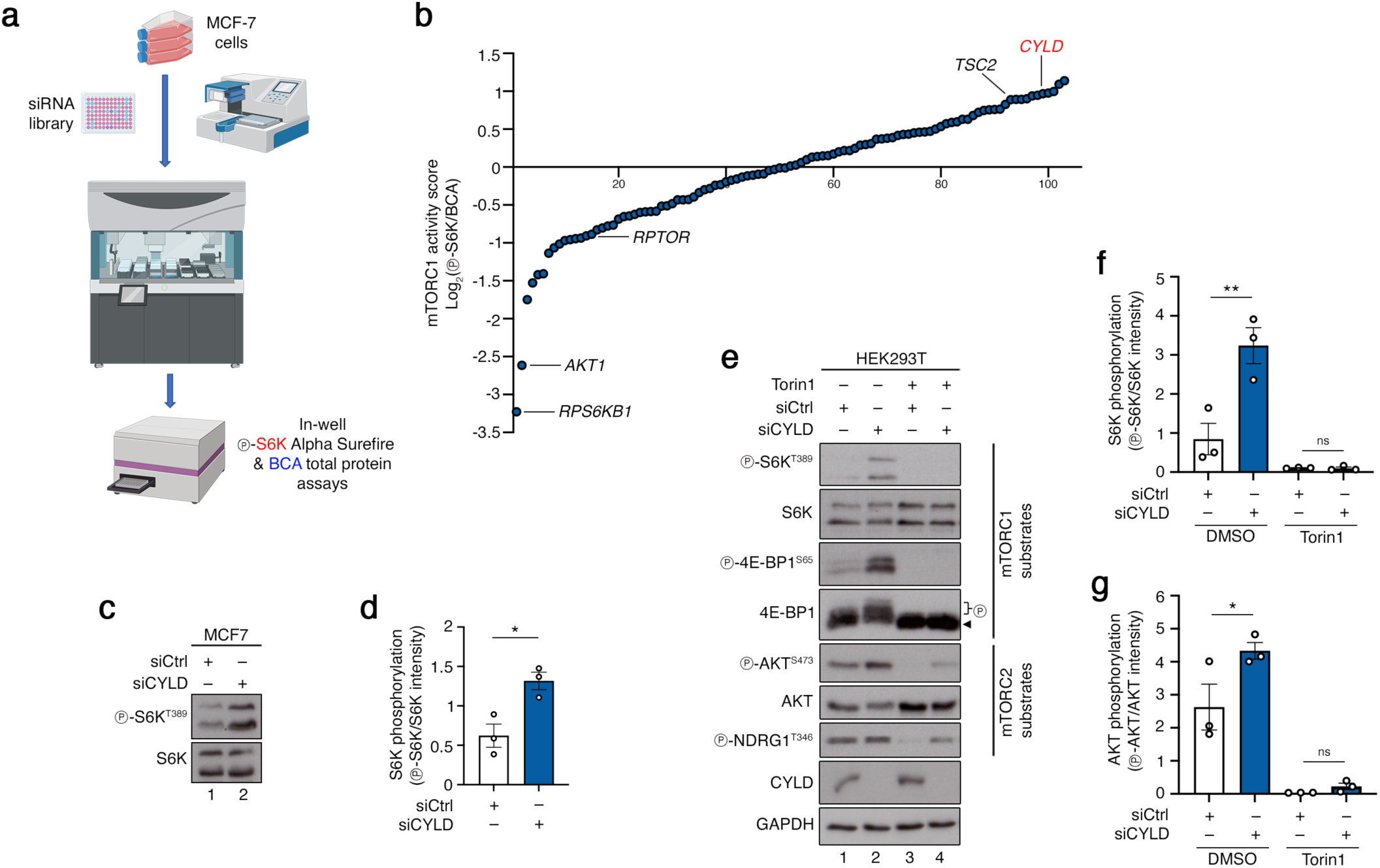
RNAi screen for DUB genes that control mTORC1 activity uncovers CYLD as a negative regulator of mTOR. **(a)** Schematic representation of the robotics-assisted, siRNA-based screen for putative mTORC1 regulators in MCF-7 cells. Levels of S6K phosphorylation normalized to total protein content was used as a proxy for mTORC1 activity. See *Methods* for more details. **(b)** Knockdown of the *CYLD* DUB gene hyperactivates mTORC1 similarly to *TSC2* knockdown in an unbiased RNAi screen. A selection of 98 human genes encoding for DUB enzymes ranked by their mTORC1 activity score in the RNAi screen shown in (a). Knockdown of *TSC2*, *RPTOR* (RAPTOR), *AKT1*, and *RPS6KB1* (S6K) itself were included in the screen as controls. **(c-d)** Immunoblots with lysates from MCF-7 cells, transiently transfected with siRNAs targeting *CYLD* or a control RNAi duplex (siCtrl), probed with the indicated antibodies (c). Quantification of S6K phosphorylation in (d). n = 3 independent experiments. **(e-g)** Immunoblots with lysates from HEK293T cells, transiently transfected with siRNAs targeting *CYLD* or a control RNAi duplex (siCtrl), treated with Torin1 for 1 h before lysis as shown, probed with the indicated antibodies (e). Quantification of S6K and AKT phosphorylation in (f) and (g), respectively. n = 3 independent experiments. Data in (d), (f), (g) shown as mean ± SEM. * p<0.05, ** p<0.01, ns: non-significant. See also Figures S1 and S2.

In confirmation of our screen data, the effect of CYLD on S6K phosphorylation was reproducible in independent knockdown experiments in MCF-7 cells (Fig. 1c,d) and human embryonic kidney HEK293T cells (Fig. 1e,f). Furthermore, CYLD knockdown increased the phosphorylation, not only of S6K, but also of another mTORC1 substrate, namely 4E-BP1 (Eukaryotic Translation Initiation Factor 4E Binding Protein 1) (Fig. 1e). In addition, the phosphorylation of both proteins was diminished upon short treatment with the catalytic mTOR inhibitor, Torin1 (Fig. 1e,f), AA starvation (Fig. S2a), glucose starvation (Fig. S2b), or growth factor removal (Fig. S2c). Collectively, these data show that CYLD acts on or upstream of mTORC1, and not directly on S6K. In addition, they indicate that the aberrantly hyperactive mTORC1 in CYLD knockdown cells can still respond to nutritional and hormonal cues. Interestingly, loss of CYLD expression led to similar increases in the phosphorylation of two downstream mTORC2 targets, namely AKT and NDRG1 (n-myc downstream regulated 1) (Fig. 1e,g). Because CYLD controls the activity of both mTORC1 and mTORC2, we speculated that CYLD may act directly on mTOR itself, as this is the catalytic subunit and a shared component of both complexes.

### CYLD negatively regulates non-degradative K63-linked ubiquitination of mTOR

As CYLD is a DUB enzyme, we next tested the potential effect of CYLD loss-of function on mTOR ubiquitination. By immunopurifying exogenously-expressed, FLAG-tagged mTOR from HEK293T cells, we could readily detect its ubiquitination by immunoblotting. Importantly, CYLD knockdown consistently and robustly increased mTOR ubiquitination further (Fig. 2a,b). Similar results were obtained with CYLD KO HEK293T cells (Fig. 2c,d), HEK293T cells overexpressing a dominant negative CYLD mutant (CYLD^C601S^) (Fig. 2e,f), and in genetically-modified mouse embryonic fibroblasts (MEFs) that express a truncated version of CYLD lacking its C-terminal DUB catalytic domain (CYLD^Δ932^) (Fig. 2g,h).

**Figure 2.**
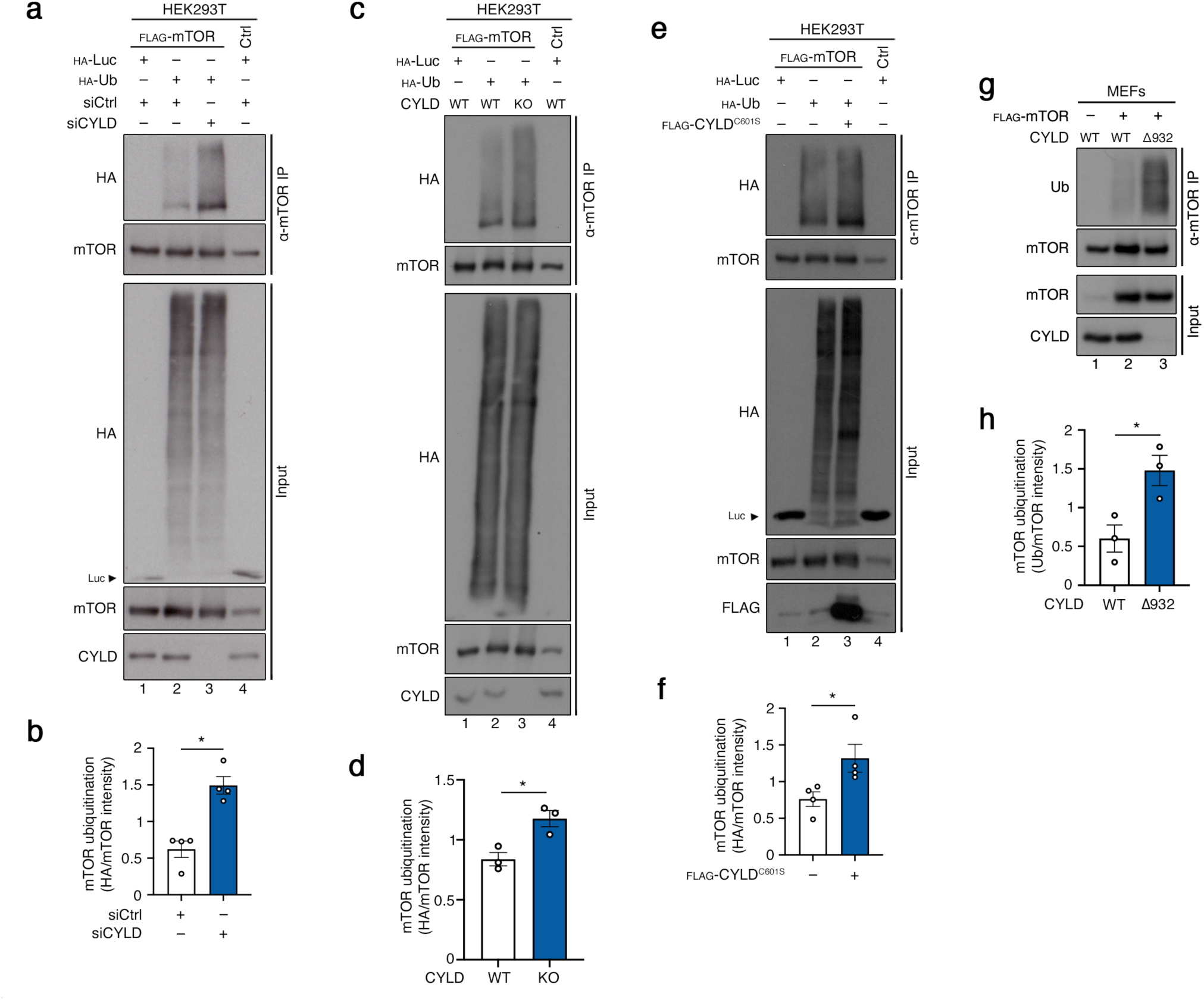
CYLD negatively regulates mTOR ubiquitination. **(a-b)** Analysis of mTOR ubiquitination in HEK293T cells transiently expressing FLAG- tagged mTOR and HA-tagged Ub, transfected with siRNAs targeting *CYLD* or a control RNAi duplex (siCtrl). Ubiquitination of immunopurified mTOR was assessed by immunoblotting with the indicated antibodies (a). Quantification of mTOR ubiquitination in (b). n = 4 independent experiments. **(c-d)** Analysis of mTOR ubiquitination in WT or CYLD KO HEK293T cells transiently expressing FLAG-tagged mTOR and HA-tagged Ub. Ubiquitination of immunopurified mTOR was assessed by immunoblotting with the indicated antibodies (c). Quantification of mTOR ubiquitination in (d). n = 3 independent experiments. **(e-f)** Analysis of mTOR ubiquitination in HEK293T cells transiently expressing FLAG- tagged mTOR, HA-tagged Ub, and a dominant-negative CYLD mutant (CYLD^C601S^) as shown. Ubiquitination of immunopurified mTOR was assessed by immunoblotting with the indicated antibodies (a). Quantification of mTOR ubiquitination in (f). n = 4 independent experiments. **(g-h)** Analysis of mTOR ubiquitination in WT or *Cyld^Δ932^* mutant MEFs stably expressing FLAG-tagged mTOR upon induction with doxycycline (250 ng/ml, 4 h before lysis) as shown. Ubiquitination of immunopurified mTOR was assessed by immunoblotting with the indicated antibodies (g). Quantification of mTOR ubiquitination in (h). n = 3 independent experiments. Data in graphs shown as mean ± SEM. * p<0.05. See also Figure S3.

Through its DUB catalytic activity, CYLD selectively removes K63- or M1-linked Ub chains from its substrates, while exhibiting very little activity towards K48-linked chains ^26, 34, 35^. As mTOR was previously shown to be decorated both by K48- and K63-linked Ub chains ^20, 36^, we next tested which type of Ub chains CYLD removes from mTOR. In either control or CYLD knockdown cells, mTOR ubiquitination was strongly increased in cells expressing K48R mutant Ub (a mutant that cannot form K48-linked chains), and much weaker in cells expressing K63R-mutant Ub (a mutant that cannot form K63-linked chains), compared to WT-Ub-expressing cells (Fig. S3a,b), indicating that mTOR is modified primarily by K63-linked Ub chains. Furthermore, while CYLD knockdown further boosted the ubiquitination of mTOR in cells expressing WT or K48R mutant Ub, it had no effect in cells expressing K63R mutant Ub (Fig. S3a). Finally, transient depletion of CYLD expression did not consistently affect the protein levels of mTOR or any other mTORC1/2 pathway component tested, with the exception of RHEB that was mildly downregulated in CYLD knockdown cells (Fig. S3c). Nevertheless, as RHEB is essential for the activation of mTORC1, lower RHEB levels cannot explain the mTORC1 hyperactivation phenotype that we observe in CYLD loss-of-function cells, suggesting the existence of a different mechanistic explanation for this effect. Taken together, our data indicate that CYLD is responsible for the removal of non-degradative, K63-linked Ub chains from mTOR.

### CYLD interacts with and deubiquitinates mTOR directly

Signals from nutrient and growth factor availability are conveyed to mTORC1 via distinct Rag GTPase heterodimers and the TSC complex, respectively, while intense interplay between these two signaling branches is also evident in mammalian cells ^14, 37–43^. Therefore, we next tested if CYLD regulates mTOR ubiquitination and activity indirectly, through these upstream signaling hubs. Interestingly, CYLD knockdown resulted in hyperactivation of mTORC1 and caused increased mTOR ubiquitination also in cells lacking expression of RagA/B (Fig. 3a-d) or the TSC1 subunit of the TSC complex (Fig. 3e-h), showing that CYLD acts independently and downstream of these regulatory complexes, possibly by acting directly on mTORC1.

**Figure 3.**
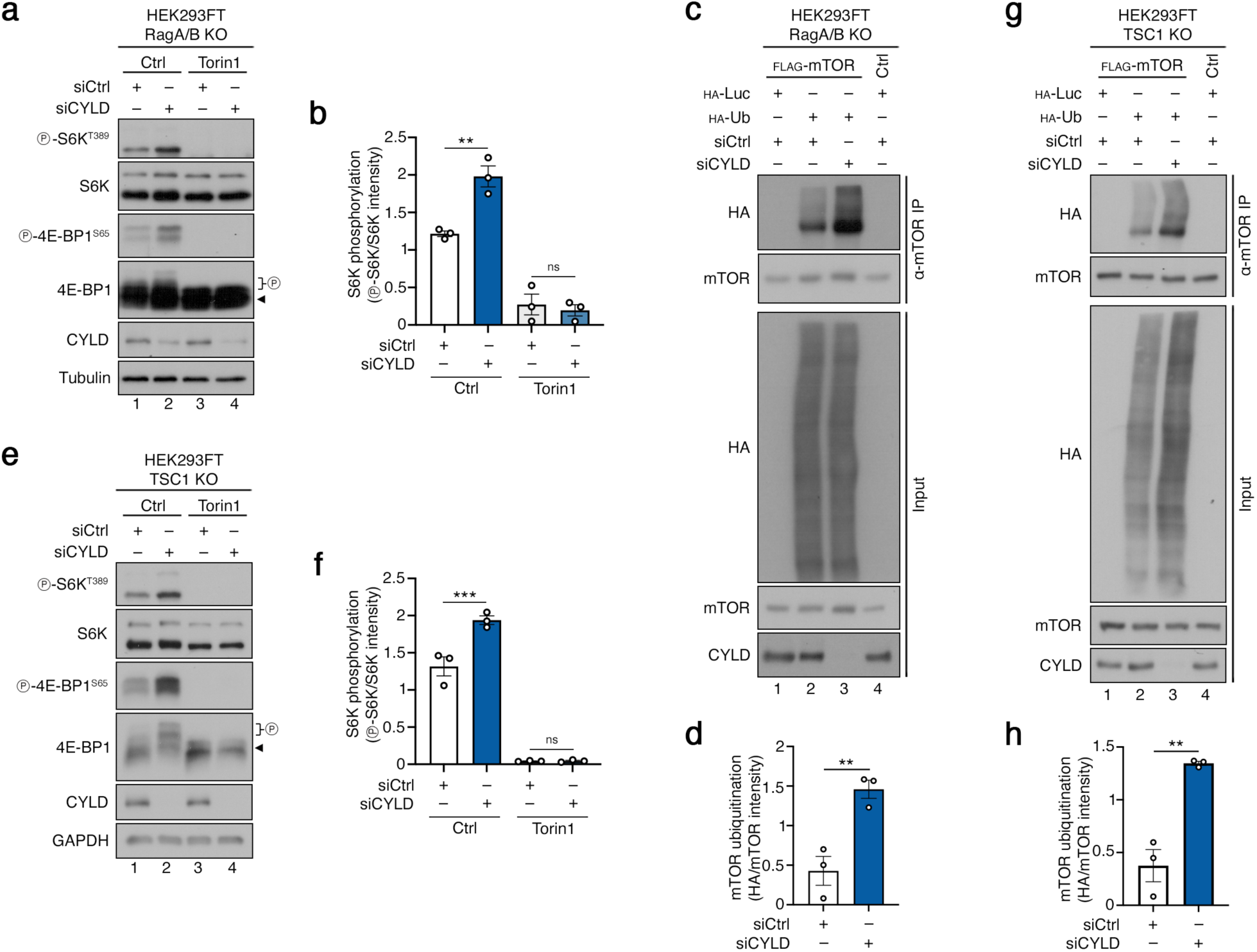
CYLD negatively controls mTORC1 activity and mTOR ubiquitination independently of key upstream mTORC1 regulators. **(a-b**) CYLD knockdown boosts mTORC1 activity also in RagA/B KO cells. Immunoblots with lysates from RagA/B KO HEK293FT cells, transiently transfected with siRNAs targeting *CYLD* or a control RNAi duplex (siCtrl), treated with Torin1 for 1 h before lysis as shown, probed with the indicated antibodies (a). Quantification of S6K phosphorylation in (b). n = 3 independent experiments. **(c-d)** CYLD knockdown boosts mTOR ubiquitination also in RagA/B KO cells. Analysis of mTOR ubiquitination in RagA/B KO HEK293FT cells transiently expressing FLAG-tagged mTOR and HA-tagged Ub, transfected with siRNAs targeting *CYLD* or a control RNAi duplex (siCtrl). Ubiquitination of immunopurified mTOR was assessed by immunoblotting with the indicated antibodies (c). Quantification of mTOR ubiquitination in (d). n = 3 independent experiments. **(e-f**) As in (a-b), but in TSC1 KO HEK293FT cells. n = 3 independent experiments. **(g-h)** As in (c-d), but in TSC1 KO HEK293FT cells. n = 3 independent experiments. Data in graphs shown as mean ± SEM. ** p<0.01, *** p<0.001, ns: non-significant.

In its canonical role, CYLD acts as a negative regulator of innate immune signaling, upstream of the IKK complex and NF-κB activation ^26^. Because IKKβ, a core kinase in TNFα/NF-κB signaling, has been shown to promote mTORC1 activation via an inhibitory phosphorylation of TSC1 ^44^, we next tested whether CYLD regulates mTOR ubiquitination via IKKβ. Although silencing of IKKβ and CYLD expression, alone or in combination, had the expected effects on NF-κB activation (Fig. S4a), IKKβ knockdown did not influence the basal or CYLD-knockdown-induced ubiquitination of mTOR (Fig. S4b).

In our experiments analyzing the ubiquitination status of mTOR, we intentionally lysed cells and immunopurified mTOR using buffers containing the detergent Triton X-100, which disrupts interactions between mTOR and most of its binding partners, including other mTORC1 components like RAPTOR ^45, 46^. Accordingly, similar experiments assessing the ubiquitination of His-tagged mTOR, purified from cells under high stringency conditions (i.e., with buffers containing 8M urea), further confirmed that CYLD knockdown boosts the ubiquitination of mTOR itself and not of any of its interaction partners (Fig. 4a,b). Moreover, recombinant CYLD protein was able to deubiquitinate immunopurified mTOR *in vitro* (Fig. 4c,d). In agreement with mTOR being a direct substrate of CYLD, we also detected reciprocal binding between the two proteins in co-immunoprecipitation (co-IP) (Fig. 4e), streptavidin pulldown (Fig. 4f), and proximity biotinylation assays (Fig. S4c).

**Figure 4.**
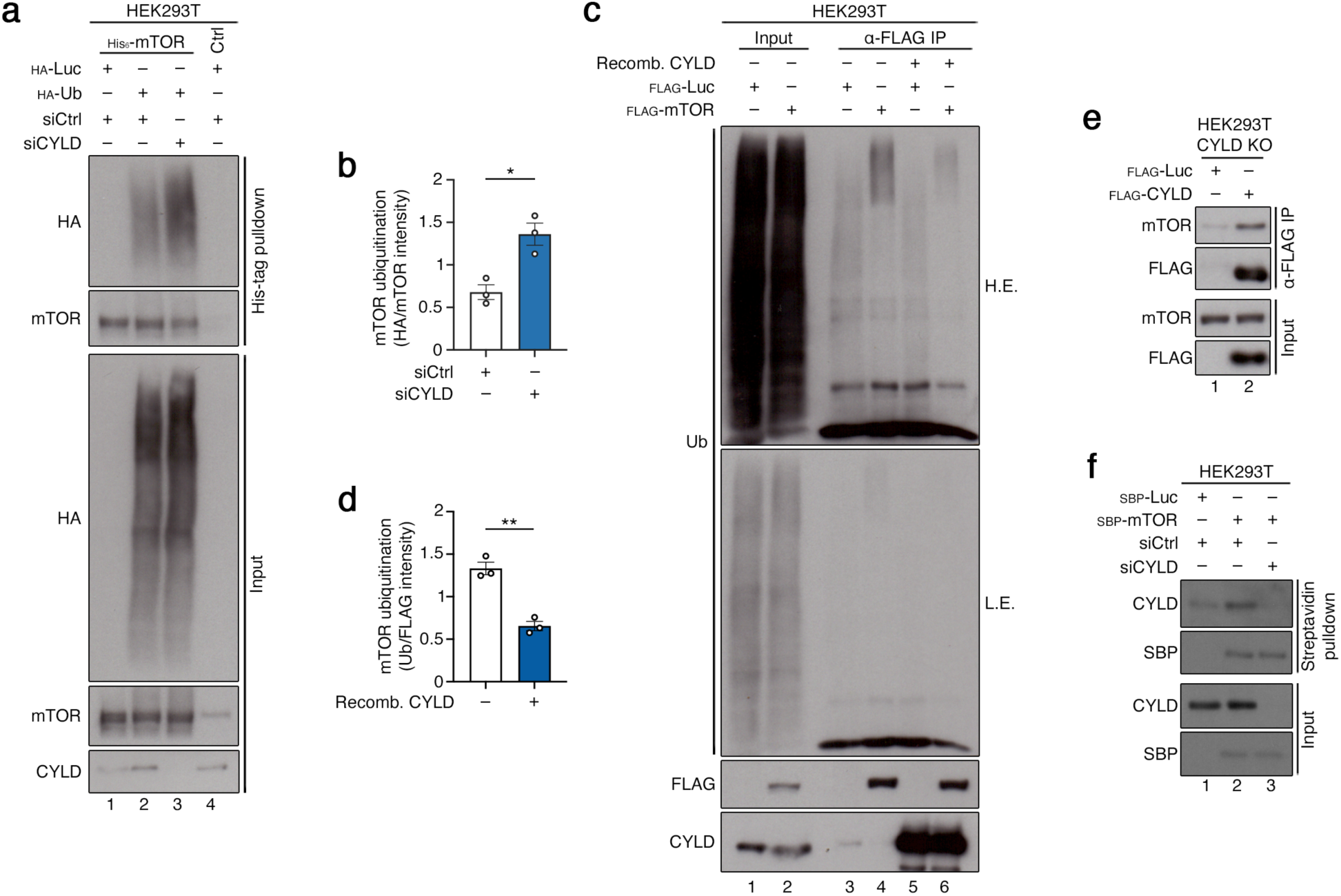
CYLD acts directly on mTOR. **(a-b)** Analysis of mTOR ubiquitination in HEK293T cells transiently expressing His6- tagged mTOR and HA-tagged Ub, transfected with siRNAs targeting *CYLD* or a control RNAi duplex (siCtrl). mTOR was purified by His-tag pulldown under stringent lysis and washing conditions to remove interacting proteins. Ubiquitination of mTOR was assessed by immunoblotting with the indicated antibodies (a). Quantification of mTOR ubiquitination in (b). n = 3 independent experiments. **(c-d)** CYLD deubiquitinates mTOR also *in vitro*. Analysis of mTOR ubiquitination in HEK293T cells transiently expressing FLAG-tagged mTOR (or an unrelated protein as control). FLAG IP samples were incubated with recombinant CYLD protein (200 ng) for 1 h before boiling as shown, and ubiquitination of immunopurified mTOR was assessed by immunoblotting with the indicated antibodies (c). Quantification of mTOR ubiquitination in (d). n = 3 independent experiments. **(e)** Exogenously-expressed CYLD interacts with endogenous mTOR. Immunoblots with FLAG IP samples and whole cell lysates (input) from CYLD KO HEK293T cells transiently expressing FLAG-tagged CYLD (or an unrelated protein as control), probed with the indicated antibodies. n = 3 independent experiments. **(f)** Exogenously-expressed mTOR interacts with endogenous CYLD. Immunoblots with streptavidin pulldown samples and whole cell lysates (input) from HEK293T cells, transiently expressing SBP-tagged mTOR (or an unrelated protein as control), transfected with siRNAs targeting *CYLD* or a control RNAi duplex (siCtrl), probed with the indicated antibodies. n = 2 independent experiments. Data in graphs shown as mean ± SEM. * p<0.05, ** p<0.01. See also Figure S4.

### CYLD regulates multi-site ubiquitination of mTOR

The human mTOR reference protein sequence (UniProt #P42345) contains 130 lysine residues. According to the PhosphoSitePlus database (www.phosphosite.org) ^47^, 47 of those lysines have been identified as ubiquitinated in previous proteomics-based studies. To test whether CYLD removes Ub chains from a single or multiple of those mTOR residues, we generated a number of mutants (mTOR^T631G^, mTOR^3xKR^, and mTOR^K2066R^) and tested their effect on mTOR ubiquitination. The mTOR^T631G^ mutant was previously described to strongly reduce the interaction between mTOR and the FBXW7 E3 ubiquitin ligase that ubiquitinates and targets mTOR for degradation ^19^. The mTOR^3xKR^ mutant (K777/782/784R) prevents the TRAF6-mediated poly- ubiquitination and activation of mTOR in response to AA stimulation ^20^. Finally, the Parkin or FBXO22 E3 ligases were previously described to ubiquitinate mTOR on K2066 to regulate its activity in response to mitochondrial stress or AA starvation, respectively ^21, 22^. In agreement with these previous studies, mTOR ubiquitination was reduced in each of the three mutants tested, compared to wild-type mTOR (Fig. 5a,b). Intriguingly, also the elevated ubiquitination of wild-type mTOR, caused by expression of dominant negative CYLD, was blunted in the mTOR mutants (Fig. 5c,d), indicating that multiple mTOR sites are likely targeted by CYLD for deubiquitination.

**Figure 5.**
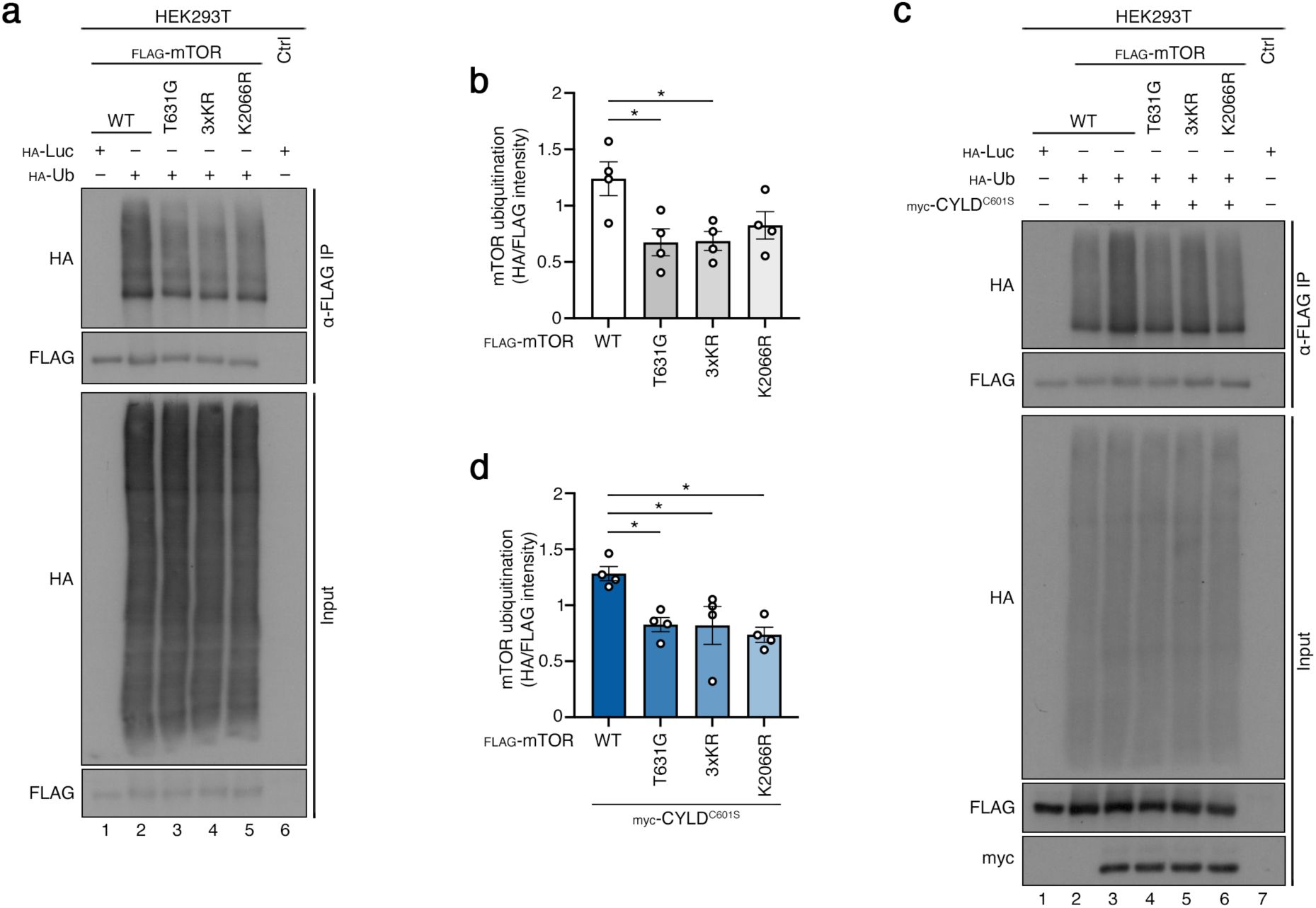
CYLD regulates multi-site ubiquitination of mTOR. **(a-b)** Multiple mTOR residues contribute to its ubiquitination under basal conditions. Analysis of mTOR ubiquitination in HEK293T cells transiently expressing FLAG- tagged WT or mutant (T631G, 3xKR, or K2066R) mTOR and HA-tagged Ub. Ubiquitination of immunopurified mTOR was assessed by immunoblotting with the indicated antibodies (a). Quantification of mTOR ubiquitination in (b). n = 4 independent experiments. **(c-d)** Multiple mTOR residues contribute to its elevated ubiquitination upon CYLD loss- of-function. Analysis of mTOR ubiquitination in HEK293T cells transiently expressing FLAG-tagged WT or mutant (T631G, 3xKR, or K2066R) mTOR, HA-tagged Ub, and a dominant-negative CYLD mutant (CYLD^C601S^) as shown. Ubiquitination of immunopurified mTOR was assessed by immunoblotting with the indicated antibodies (c). Quantification of mTOR ubiquitination in (d). n = 4 independent experiments. Data in graphs shown as mean ± SEM. * p<0.05

### CYLD restricts mTOR-driven protein synthesis and cell growth

We show here that CYLD functions as a sentinel of mTORC1 hyperactivation via directly controlling its ubiquitination. Therefore, we next aimed to investigate the physiological role of mTOR’s regulation by CYLD. Given the well-established role of mTORC1 as a master regulator of protein synthesis and cell growth via the phosphorylation of S6K and 4E-BP1, we assessed *de novo* protein synthesis by using a puromycin incorporation assay in two distinct CYLD loss-of-function cellular models. Consistent with S6K and 4E-BP1 phosphorylation being robustly upregulated in cells with reduced CYLD expression, we observed elevated translation rates in both CYLD knockdown and knockout cells, that were strongly inhibited by treatment with Torin1 and diminished by cycloheximide (CHX) (Fig. 6a,b and Fig. S5a,b). Accordingly, CYLD knockdown led to an increase in cell size, comparable to that seen in TSC2-silenced cells, that was fully reversed by mTOR inhibition with Torin1 (Fig. 6c). Similar results were obtained when comparing the size of CYLD-knockout cells to the respective CYLD-proficient controls (Fig. S5c).

**Figure 6.**
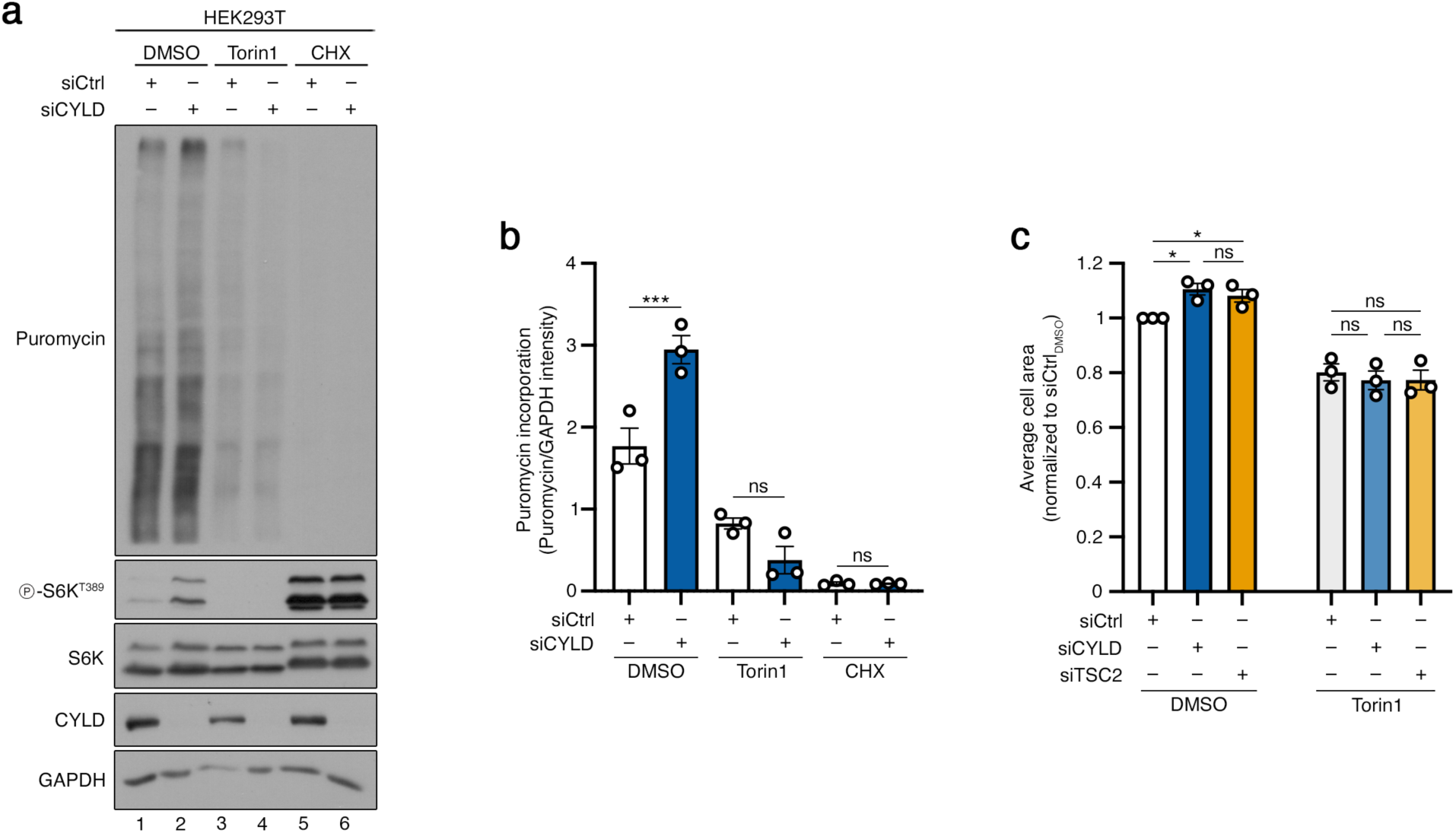
CYLD restricts mTOR-driven protein synthesis and cell growth. **(a-b)** Elevated translation rates in CYLD knockdown cells are reversed by mTOR inhibition. Puromycin incorporation assays in HEK293T cells, transiently transfected with siRNAs targeting *CYLD* or a control RNAi duplex (siCtrl), treated with Torin1 (250 nM, 16 h), CHX (100 µM, 4 h), or DMSO as control, analyzed by immunoblotting with the indicated antibodies (a). Quantification of puromycin incorporation into nascent polypeptide chains in (b). n = 3 independent experiments. **(c)** Increased cell size of CYLD knockdown cells is reversed by mTOR inhibition. Cell area measurements from HEK293T cells, transiently transfected with siRNAs targeting *CYLD*, *TSC2*, or a control RNAi duplex (siCtrl), using an IncuCyte S3 live-cell imaging and analysis system. Torin1 (250 nM, 16 h) was used to inhibit mTOR. n = 3 independent experiments. Data in graphs shown as mean ± SEM. * p<0.05, *** p<0.001, ns: non-significant. See also Figures S5 and S6.

In addition to its effects on mTORC1 activity, silencing of CYLD also caused elevated phosphorylation of mTORC2 substrates that persisted even in serum-starved cells (Fig. S2c and Fig. S6). As mTORC2 promotes survival, in part via the phosphorylation of AKT, downregulation of its activity upon prolonged serum starvation in control cells caused the activation of cell-death-related pathways, as evidenced by the induction of PARP1 cleavage in immunoblotting experiments (Fig. S6, lanes 1 and 3). In contrast, the persistent mTORC2 activation in growth-factor-deprived cells upon CYLD- knockdown was associated with compromised activation of cell death pathways (Fig. S6, lanes 3 and 4), while staurosporine, a drug that induces cell death independently of mTORC2, induced PARP1 cleavage to a similar extent in control and CYLD knockdown cells (Fig. S6, lanes 5 and 6). Taken together, our data show that CYLD restricts protein synthesis and cell growth, and mediates the serum-starvation-induced cell death via monitoring mTORC1 and mTORC2 activities.

### CYLD regulates *C. elegans* ageing via suppressing mTOR activity

mTOR plays a central role in the ageing process and dysregulation of its activity has been linked to cellular dysfunction in aged tissues ^12^. Therefore, we tested the role of *cyld-1*, the *C. elegans* CYLD ortholog, in the regulation of mTOR activation and ageing in worms. In *C. elegans*, depletion of mTOR signaling through loss-of-function mutations in *raga-1* (the RagA ortholog) dramatically extends lifespan ^48, 49^. Strikingly, silencing of *cyld-1* fully reversed the lifespan extension observed in *raga-1* mutant worms, suggesting that *cyld-1* activates mTOR downstream of *raga-1* (Fig. 7a). Indeed, using the nuclear localization of a fluorescent HLH-30/TFEB reporter as a readout for mTOR activity, we observed lower nuclear presence of HLH-30 upon *cyld- 1* knockdown in *raga-1* mutant worms (Fig. 7b,c). These data indicate that CYLD maintains mTORC1 activity under control to regulate key physiological processes, not only in cultured cells, but also at the organismal level.

**Figure 7.**
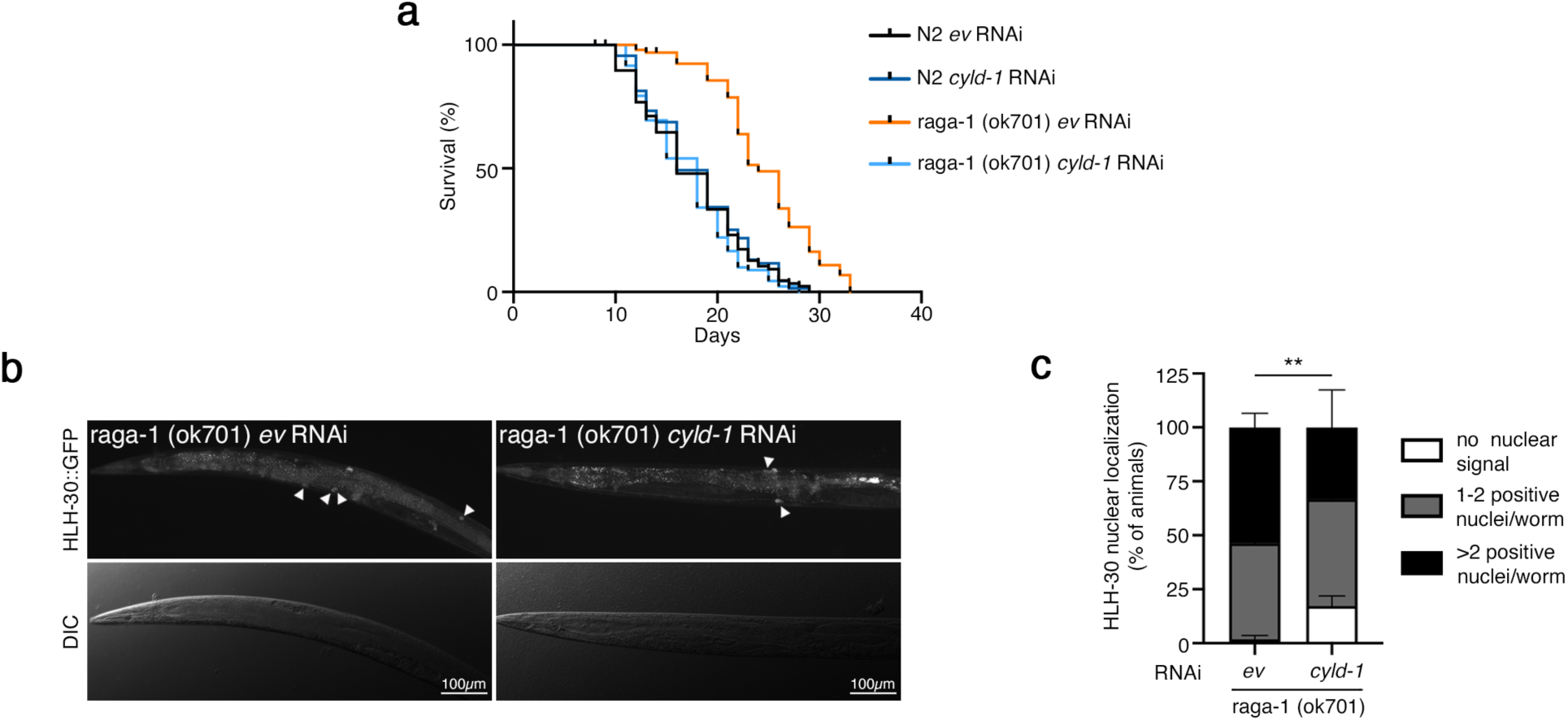
*cyld-1* is required for the lifespan extension and the nuclear HLH- 30/TFEB localization observed in *raga-1* mutants. **(a)** Knockdown of *cyld-1* after development reverses the extended longevity induced by the loss-of-function mutation in *raga-1*. Lifespan analysis of wild-type (N2) and *raga-1* mutant *C. elegans* with or without knockdown of *cyld-1*. n = 3 independent experiments **(b-c)** Knockdown of *cyld-1* blunts the nuclear localization of HLH-30/TFEB, characteristic of *raga-1* mutant worms. HLH-30::GFP reporter localization and differential interference contrast (DIC) imaging in *raga-1* mutant with or without knockdown of *cyld-1*. Scale bars: 100 µm (b). Quantification of HLH-30 nuclear localization in (c). Data shown as mean ± SEM. ** p<0.01. n = 3 independent experiments

### Hyperactivation of mTORC1 in skin biopsies of cylindromatosis patients

CCS is clinically characterized by the development of multiple tumors of the skin appendages, mainly in the head and neck region, including cylindromas, spiradenomas, and trichoepitheliomas. These skin lesions are generally of epidermal origin, which is in agreement with CYLD showing the highest expression in suprabasal keratinocytes among the different skin cell types ^50, 51^. The role of CYLD inactivation in cylindromatosis is generally attributed to the constitutive activation of pro-survival and anti-apoptosis-related signaling, including the NF-κB and JNK pathways ^24–26, 52^. As a key regulator of virtually all major cellular processes, dysregulation of mTOR activity plays a central role in cancer development and growth. Therefore, we sought to investigate the activation state of mTORC1 in skin biopsies of patients that have been clinically diagnosed with cylindromatosis as well as based on histopathological and/or genetic testing. Although cylindromatosis is a rare genetic syndrome, we identified two distinct individuals. Patient #1 harbors a mutation in the *CYLD* locus that introduces a premature termination codon (CYLD c.2797C/T, p.Arg933Ter), resulting in a catalytically-inactive form of the protein, further confirming the presence of CCS/BSS. The pathological evaluation of these skin lesions showed tumors of mixed phenotype, with both spiradenoma- and cylindroma-like structures, regressive changes, and cystic and hyaline degeneration. Lesions excised from patient #2 showed a similar histopathological phenotype, typical of CCS, with a higher prevalence of cylindromas (Fig. 8a).

**Figure 8.**
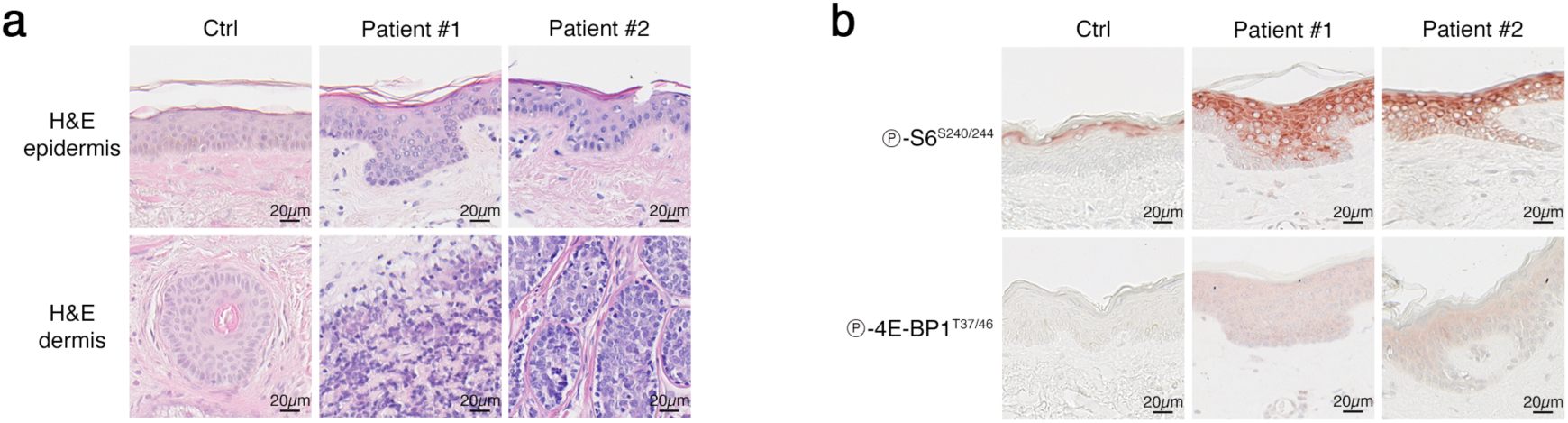
Aberrantly elevated mTORC1 activity in skin biopsies from CCS/BSS patients. **(a)** Representative hematoxylin and eosin (H&E) staining of epidermal and dermal areas from normal skin (Ctrl) and skin biopsies from two independent CCS patients. Scale bars: 20 µm. **(b)** Representative immunohistochemistry staining of epidermal areas from normal skin (Ctrl) and skin biopsies from two independent CCS patients. Phosphorylation of S6 and 4E-BP1 were used as readouts for mTORC1 activity. Scale bars: 20 µm.

In the epidermis of normal skin, high mTORC1 activity (most commonly using S6 or 4E-BP1 phosphorylation as readouts) is primarily detected in the granular (stratum granulosum) and spinous (stratum spinosum) layers ^53^, as well as in hair follicles and various glands ^53, 54^. Accordingly, we could detect moderate phosphorylation of S6 in the cells comprising the upper layer of the epidermis, just below the stratum corneum (Fig. 8b). In contrast, 4E-BP1 phosphorylation was almost undetectable in normal skin samples (Fig. 8b). Notably, the phosphorylation of both S6 and 4E-BP1 was dramatically increased in the epidermis of both cylindromatosis patients (Fig. 8b). Moreover, we observed an expansion of the phopsho-S6-positive area, as well as mild epidermal hyperplasia (acanthosis) with increased thickness and average cell size in the stratum spinosum of the patients’ skin (Fig. 8b). In sum, the analysis of skin biopsies from CCS patients reveals that CYLD loss-of-function is associated with aberrant hyperactivation of mTORC1 signaling, similarly to our observations in cell culture models of cylindromatosis.

## Discussion

The data presented here reveal CYLD as a DUB enzyme that acts directly on mTOR to constrain its activity. Our findings are consistent with previous observations hinting at the existence of potential links between the two proteins: for instance, the phosphorylation of S6K, and of its downstream target S6, were found to be elevated in hippocampus samples from CYLD-deficient mice. In addition, CYLD was reported to interact with mTOR in wild-type mouse hippocampus homogenates ^55^. Accordingly, overexpression of CYLD in cardiomyocytes of transgenic mice lowered S6K phosphorylation ^56^. Together with our work, these reports support a negative role for CYLD in the regulation of mTORC1 activity in diverse cell types and species.

Non-degradative ubiquitination of client proteins can alter their function by diverse ways: it can form new interaction surfaces that promote binding of the ubiquitinated protein to upstream regulators and/or downstream effectors and substrates. One such example is the polyubiquitination of RIPK1 that was shown to facilitate the recruitment of the TAB2 (TAK1 binding protein 2) and TAB3 adaptor proteins via their interaction with K63-linked Ub chains on RIPK1, a recruitment that is required for the activation of downstream kinases like TAK1 and the IKK complex ^57^. However, as CYLD regulates the ubiquitination of mTOR in the context of both mTORC1 and mTORC2, and because these two complexes have distinct sets of upstream regulators and downstream targets, we find it less likely that CYLD controls mTOR activity via a similar mechanism. Alternatively, ubiquitination has been described to reversibly regulate enzymatic activity by inducing conformational changes to the target protein ^58^. Our data indicate that CYLD is responsible for the removal of K63-linked Ub chains from multiple residues on mTOR. In fact, the sites that we tested in our mTOR ubiquitination assays are not concentrated at a particular domain of mTOR, but are rather spread over different regions of the protein (e.g., the T631 and K777/782/784 sites are located in the HEAT repeat domain, whereas K2066 is in the kinase domain). This suggests that CYLD-mediated deubiquitination of mTOR may be affecting multiple aspects of the mTOR activation process that collectively result in keeping its activity under tight control. Future work will be necessary to disentangle the precise molecular effects that the addition of non-degradative ubiquitin chains has on mTOR conformation, dimerization, or interaction with its regulators and effector proteins; and to understand the specific role of CYLD in all of these processes.

Inactivating mutations in *CYLD* are causative of CCS, a rare human disease that manifests with tumors of the skin appendages ^30^. Along these lines, genetic inactivation of CYLD specifically in the epidermis sensitizes mice to the development of various skin tumor types upon topical challenge with carcinogens ^59^. Of note, both mTOR complexes are known to play key roles in cancer growth and development in the skin ^60^; and are also essential for skin differentiation and homeostasis ^54^. Here, we report high mTORC1 activity primarily in the stratum granulosum and spinosum layers of the epidermis of cylindromatosis patients, as also shown previously for healthy individuals ^53, 54^. A potentially interesting coincidence is the expression pattern of CYLD in the skin, showing highest expression in suprabasal keratinocytes ^50, 51^, that closely matches the pattern of high mTORC1 signal in cylindromatosis patient samples. Based on this, it is plausible to speculate that high CYLD expression in the upper layers of the epidermis normally keeps mTORC1 activity under close surveillance, which is why CYLD deficiency in CCS results in the strongest dysregulation of mTORC1 in this area.

Besides inactivating mutations in its gene locus, CYLD expression was shown to be downregulated, or even lost, in various human colon and hepatocellular carcinomas^61^. In light of our findings, this transcriptional repression of CYLD may be contributing to tumor development and progression through the upregulation of TOR signaling (in addition to the classical anti-apoptotic NF-κB and JNK signaling pathways). Further supporting the hypothesis that mTOR dysregulation may be contributing to the symptomatology of cylindromatosis, previous reports have noted that affected patients exhibit clinical features similar to those of tuberous sclerosis patients—a distinct genetic disorder characterized by the loss of TSC function and hyperactivation of mTOR signaling ^62^.

Using the *C. elegans* model to study the role of the CYLD-mTOR axis in ageing, we found that the lifespan extension of low-TOR-activity mutants can be reversed by CYLD knockdown in worms. Interestingly, mice expressing a dominant negative CYLD mutant (CYLD^C601S^) under the control of keratin 5 promoter show signs of premature ageing in the skin and in multiple other tissues. Also, middle-aged or old mutant mice spontaneously develop tumors in many of their organs ^63^. Although the role of mTOR dysregulation in this setting has not been investigated so far, these studies suggest that the role of CYLD in ageing is likely evolutionarily conserved, at least from worms to mammals.

Taken together, our findings highlight CYLD as a sentinel of mTORC1 hyperactivation via directly controlling its ubiquitination, while CYLD loss-of-function is associated with dysregulated mTOR signaling. Due to the emerging potential role of the CYLD-mTOR axis in skin tumor development, we put forward the hypothesis that pharmacological inhibition of mTOR may constitute a promising therapeutic approach to treat cylindromatosis patients in the future.

## Methods

### Cell culture

All cell lines were grown at 37 °C, 5% CO_2_. Human female embryonic kidney HEK293FT cells (Research Resource Identifier (RRID): CVCL_6911) and HEK293T cells (RRID: CVCL_0063), human female breast adenocarcinoma MCF-7 cells (HTB- 22, ATCC; RRID: CVCL_0031) and immortalized mouse embryonic fibroblasts (MEFs), were cultured in high-glucose Dulbecco’s Modified Eagle Medium (DMEM) (#41965-039, Gibco), supplemented with 10% fetal bovine serum (FBS) (#F7524, Sigma; #S1810, Biowest). Culture media of MCF-7 cells were also supplemented with 1x non-essential amino acids (#11140-035, Gibco). All media were supplemented with 1x penicillin– streptomycin (#15140-122, Gibco).

HEK293FT cells were purchased from Invitrogen (#R70007). The wild-type control and CYLD^Δ932^ immortalized MEFs, harboring the R932X mutation, were generated in the Pasparakis lab (described in ^64^). The identity of the HEK293FT and MCF7 cells was validated by the Multiplex human Cell Line Authentication test (Multiplexion GmbH), which uses a single nucleotide polymorphism (SNP) typing approach, and was performed as described at www.multiplexion.de. No commonly misidentified cell lines were used in this study. All cell lines were regularly tested for *Mycoplasma* contamination, using a PCR-based approach and were confirmed to be *Mycoplasma*- free.

### Cell culture treatments

Amino acid (AA) starvation experiments were performed as described previously ^39, 40^. In brief, custom-made starvation medium was formulated according to the Gibco recipe for high-glucose DMEM, specifically omitting all amino acids. The medium was filtered through a 0.22-μm filter device and tested for proper pH and osmolality before use. For the respective AA-replete (+AA) treatment media, commercially available high-glucose DMEM was used. All treatment media were supplemented with 10% dialyzed FBS (dFBS) and 1x penicillin-streptomycin. For this purpose, FBS was dialyzed against 1x phosphate-buffered saline (PBS) through 3.500 MWCO dialysis tubing. For basal (+AA) conditions, the culture media were replaced by +AA treatment media one hour before lysis. For AA starvation (–AA), culture media were replaced by starvation media for one hour. For glucose starvation experiments, cells were cultured for one hour in glucose-free DMEM (#11966025, Gibco) supplemented with 10% dFBS and 1x penicillin-streptomycin. For the respective control wells, the culture media were replaced by high-glucose DMEM containing 10% dFBS and 1x penicillin-streptomycin at the beginning of the experiment. For growth factor starvation experiments, complete culture media were replaced by FBS-free DMEM supplemented with 1x penicillin-streptomycin for 16 hours (for mTOR activity assays) or for 24 hours (for PARP1 cleavage assays). For control wells, media were exchanged will fresh full media 16 or 24 hours, respectively, prior to lysis.

To inhibit mTOR kinase activity, Torin1 (#14379, Cell Signaling Technology) was added directly in the culture media (final concentration 250 nM) for one hour (for mTOR activity assays), for 16 hours (for puromycin incorporation assays), or for 72 hours (for cell size measurements). To inhibit protein synthesis, cycloheximide (#239763, Sigma) was added to the media (final concentration 100 µM) for four hours. To induce the activation of cell death pathways, Staurosporine (#569397, Sigma) was added to the culture media (final concentration 1 µM) for six hours. DMSO (#4720.1, Roth) was used as control for all treatments.

### Antibodies

A list of all primary antibodies used in this study is found in Table S2.

### Plasmids and molecular cloning

The luciferase-based NF-κB reporter vector (3xKBL) was described in ^65^, while the internal control pRL-null vector was purchased by Promega (#PAE2271, Promega). The pRK5-HA-Ubiquitin-WT (Addgene plasmid #17608; deposited by the Ted Dawson lab) ^66^ and the pcDNA3-FLAG-mTOR (Addgene plasmid #26603; deposited by the Jie Chen lab) ^67^ plasmids were obtained from Addgene. The K48R and K63R ubiquitin mutants were generated by site-directed mutagenesis using appropriate oligos, and the resulting PCR products were cloned into the EcoRI and Acc65I restriction sites of the pRK5-HA-Ubiquitin-WT vector. The pcDNA3-His_6_-mTOR and pcDNA3-HA-mTOR expression vectors were generated by replacing the N-terminal FLAG tag in the pcDNA3-FLAG-mTOR vector with His_6_ and HA tags, respectively, using appropriate oligos, and the PCR products were cloned into the HindIII and NheI restriction sites of the original pcDNA3-FLAG-mTOR.

The pITR-TTP-FLAG-APEX2-mTOR was generated by PCR amplification of the APEX2 sequence from the pITR-TTP-hGRASP55-APEX2-myc-His_6_ plasmid ^68^ using oligonucleotides that also contained the FLAG tag sequence, and inserted into the SfiI/NheI sites of the pITR-TTP1 backbone. Next, mTOR was PCR amplified from the pcDNA3-FLAG-mTOR plasmid and cloned into the NotI/XhoI sites of the pITR-TTP- FLAG-APEX2 vector. To generate the pITR-TTP-FLAG-mTOR expression vector, the FLAG-APEX2 sequence was removed using the SfiI/NotI restriction sites, and the FLAG tag sequence was reintroduced into the same sites, in-frame with mTOR, using a double-stranded DNA oligo.

The pcDNA3-FLAG-CYLD and pcDNA3-FLAG-CYLD-C601S plasmids were described in ^26^, while the pcDNA4/TO/SBP-mTOR was described in ^69^. The pcDNA3- myc-CYLD-C601S, pcDNA3-FLAG-mTOR-T631G, pcDNA3-FLAG-mTOR-3xKR (K777/782/784R) and pcDNA3-FLAG-mTOR-K2066R expression vector were generated by PCR amplification using appropriate oligos and the NEBuilder HiFi DNA Assembly Master Mix (#E2621, New England BioLabs). In brief, CYLD-C601S was amplified from the pcDNA3-FLAG-CYLD-C601S plasmid ^26^ and assembled into the EcoRI/NotI sites of pCMV-myc (#635689, Clontech). Next, myc-CYLD-C601S was subcloned into the XbaI/BamHI sites of pcDNA3. For the generation of the mTOR T631G and 3xKR mutants, GeneArt strings (Thermo Scientific) were designed to contain the desired mutations and cloned into the EcoRV/KasI sites of pcDNA4/TO/SBP-mTOR, replacing the wild-type sequence. Next, part of the mTOR sequence was PCR amplified from the pcDNA4/TO/SBP-mTOR-T631G or 3xKR plasmids and assembled into the HindIII/XbaI sites of pcDNA3-FLAG-mTOR. The mTOR-K2066R mutant was generated by site-directed mutagenesis using appropriate oligonucleotides, and the PCR product was assembled into the HindIII/XbaI sites of the pcDNA3-FLAG-mTOR vector. To generate the negative control HA-Luc and SBP- Luc expression vectors, the luciferase cDNA sequence was amplified from pcDNA3- FLAG-Luc ^39^ and cloned into the EcoRI/NotI sites of pcDNA3, or the BamHI/AgeI sites of pcDNA4/TO/SBP, respectively.

All restriction enzymes were purchased from Fermentas/Thermo Scientific. The integrity of all constructs was verified by sequencing. All DNA oligonucleotides and donor templates used in this study are listed in Table S3.

### Plasmid DNA transfections

Plasmid DNA transfections in HEK293FT and HEK293T cells were performed using Effectene transfection reagent (#301425, QIAGEN), according to the manufacturer’s instructions. MEFs were transfected using ViaFect transfection reagent (#E4981, Promega), as per the manufacturer’s instructions.

### Generation of stable cell lines

Stable cell lines expressing FLAG-tagged mTOR in WT and CYLD^Δ932^ (CYLD^R932X^) MEFs were generated using a doxycycline-inducible, sleeping-beauty-based system ^68, 70^. The pITR-TTP-FLAG-mTOR construct was transfected in a 10:1 ratio with the transposase-expressing pCMV-Trp vector. Forty-eight hours post-transfection, cells were selected with 3 μg/ml puromycin (#A11138-03, Thermo Fisher Scientific) and a polyclonal cell population was used for further experiments. Expression of mTOR was induced with 250 ng/ml of doxycycline (#D9891, Sigma) for four hours before cell lysis. For proximity biotinylation experiments, stable cell lines expressing a FLAG-APEX2- mTOR fusion in WT or CYLD KO HEK293T cells were generated using the same approach as described above and a pITR-TTP-FLAG-APEX2-mTOR plasmid construct. A polyclonal cell population was used for further experiments, and expression of FLAG-APEX2-mTOR was induced for 24 hours with 10 ng/ml doxycycline for WT cells 50 ng/ml of doxycycline for CYLD KO cells to achieve equal expression levels between genotypes.

### Gene silencing experiments

Transient knockdown of *CYLD*, *IKBKB* (ΙΚΚβ), and *TSC2* was performed using siGENOME (pool of 4) gene-specific siRNAs (Horizon Discovery). An siRNA duplex targeting the *Renilla reniformis* luciferase gene (RLuc) (#P-002070-01-50, Horizon Discovery) was used as control. Transfections were performed with 20 nM siRNA and the Lipofectamine RNAiMAX transfection reagent (#13778075, Thermo Fisher Scientific) in 12-well plates, according to the manufacturer’s instructions. Cells were harvested 72 hours post-transfection and knockdown efficiency was verified by immunoblotting or functional assays (e.g., NF-κB reporter assays; cell size measurements).

For RNAi experiments in *C. elegans*, worms were fed *E. coli* (HT115) containing an empty control vector (L4440) or expressing double-stranded RNAi when they reached adulthood. The *cyld-1* RNAi constructs were obtained from the Vidal library. The RNAi construct was sequence verified.

### Generation of knockout cell lines

The HEK293FT RagA/B KO and TSC1 KO cells were described previously ^69, 71^. The CYLD KO HEK293T cells were generated in the Mosialos lab using standard CRISPR/Cas9-based methods and the pD1301-AD (CMV-Cas9-2A-GFP, Cas9- ElecD; Horizon Discovery) all-in-one plasmid expressing Cas9-DasherGFP and the sgRNA guides. Targeting of the CYLD locus was performed using two independent sgRNAs recognizing sequences 5’-GTTAATATCACAATGAGTTC-3’ and 5’- GTATATTCAAGATCGTTCTG-3’. Single-cell clones were generated by single cell dilution and knockout clones were validated by immunoblotting, genomic DNA PCRs, and functional assays.

### RNAi screen for DUB genes as putative mTORC1 regulators

To identify DUB enzymes that may function as putative mTORC1 regulators, an unbiased RNAi screen was performed in human female breast adenocarcinoma MCF- 7 cells, using S6K phosphorylation as a readout for mTORC1 activity. In brief, a genome-wide arrayed siRNA library (pool of 4 siGENOME siRNA reagents per gene) was obtained from Horizon Discovery, and a subset corresponding to 98 DUB genes was selected for follow-up analysis. The DUBs were selected based on a database created by Zhe Xue, Yue Zhao, and Mark Knepper in the Epithelial Systems Biology Laboratory (https://esbl.nhlbi.nih.gov/) at the National Heart, Lung and Blood Institute (https://esbl.nhlbi.nih.gov/Databases/KSBP2/Targets/Lists/DUBs/). Positive and negative upstream mTORC1 pathway genes (AKT1, RAPTOR, TSC2), as well as S6K itself, were included in the screen as controls.

Transient knockdowns were performed by following a reverse transfection protocol. The DharmaFECT 1 transfection reagent (#T-2001-03, Horizon Discovery) diluted in RPMI media was dispensed directly in 384-well plates containing the siRNA reagents, followed by addition of the cell suspension (1000 cells per well in 30 μl antibiotic-free DMEM media containing 10% FBS) using a Multidrop Combi dispenser (Thermo Fisher Scientific). Three days post-transfection, the culture media were replaced with fresh DMEM containing 10% dialyzed FBS, and, 4 hours later, the media were removed completely by aspiration and cells were lysed in-well by the addition of 20μl 1x lysis buffer (#TGR703S50K, PerkinElmer) and shaking for 10 min at 350 rpm on an orbital shaker. Lysates (4 μl) were transferred to 384-well assay plates using a Biomek FXp liquid handling robotic device (Beckman Coulter). The activation state of mTORC1 was determined by measuring the phosphorylation levels of S6K (NP_003152) using the AlphaScreen SureFire p70 S6K (p-Thr229) Assay kit (#TGR703S50K, PerkinElmer) as per the manufacturer’s instructions. Proxiplates were read with an Enspire plate reader (PerkinElmer) using standard AlphaScreen settings. Total protein levels were measured in a separate assay plate using a BCA Protein Assay Kit (#23225, Pierce) according to manufacturer’s instructions. Normalized S6K phosphorylation values were calculated as the ratio of phospho-S6K to total protein signals, and the genes were ranked based on the ‘mTORC1 activity score’ that was calculated as log2- transformed phospho-S6K to total protein signal ratio (Table S1).

### Analysis of genetic alterations in cancer

Genetic alterations (mutations and copy number alterations) in the *CYLD*, *TSC2*, *TSC1* and *FLCN* genes were analyzed using the cBioPortal platform for Cancer Genomics (www.cbioportal.org) ^72, 73^. Data from the TCGA PanCancer Atlas cohort (10198 samples/patients in 32 studies) were queried to assess the presence of mutations and copy number alterations (CNAs) across multiple cancer types. Samples with alterations of unknown significance (783), and samples/patients that have not been profiled for all queried genes (769) were excluded from the analysis. Data were retrieved and visualized using cBioPortal’s built-in analysis tools (v6.0.25).

### Cell lysis and immunoblotting

For standard sodium dodecyl sulfate–polyacrylamide gel electrophoresis (SDS– PAGE) and immunoblotting experiments, cells from a well of a 12-well plate were treated as indicated in the figures, washed once with serum-free DMEM, and lysed in 250 μl of ice-cold Triton lysis buffer (50 mM Tris pH 7.5, 1% Triton X-100, 150 mM NaCl, 50 mM NaF, 2 mM Na-vanadate, 0.011 g/ml beta-glycerophosphate), supplemented with 1x PhosSTOP phosphatase inhibitors (#04906837001, Roche) and 1x cOmplete protease inhibitors (#11697498001, Roche), for 10 minutes on ice.

Samples were clarified by centrifugation (19000 g, 15 min, 4 °C) and supernatants transferred to a new tube. Protein concentration was determined using a Protein Assay Dye Reagent (#5000006, Bio-Rad). Normalized samples were boiled in 1x SDS sample buffer for 5 min at 95 °C (6x SDS sample buffer: 350 mM Tris-HCl pH 6.8, 50% glycerol, 600 mM DTT, 12.8% SDS, 0.12% bromophenol blue).

Protein samples were subjected to electrophoretic separation on SDS-PAGE and analysed by standard Western blotting techniques. In brief, proteins were transferred to nitrocellulose membranes (#10600002 or #10600001, Amersham) and stained with 0.2% Ponceau solution (#33427-01, Serva) to confirm equal loading. Membranes were blocked with 5% skim milk powder (#42590, Serva) in PBS-T [1x PBS, 0.1% Tween- 20 (#A1389, AppliChem)] for one hour at room temperature (RT), washed three times for 10 min with PBS-T and incubated with primary antibodies in PBS-T with 5% bovine serum albumin (BSA; #10735086001, Roche) rotating overnight at 4 °C. Antibody dilutions are provided in Table S2. The next day, membranes were washed three times for 10 min with PBS-T and incubated with appropriate HRP-conjugated secondary antibodies (1:10000 in PBS-T, 5% milk) for one hour at RT. Signals were detected by enhanced chemiluminescence (ECL), using the ECL Western Blotting Substrate (#W1015, Promega); or SuperSignal West Pico PLUS (#34577, Thermo Scientific) and SuperSignal West Femto Substrate (#34095, Thermo Scientific) for weaker signals. Immunoblot images were captured on films (#28906835, GE Healthcare; #4741019289, Fujifilm). Blots were quantified using GelAnalyzer 19.1.

### Immunoprecipitation and mTOR ubiquitination analysis

For immunoprecipitation experiments, cells were lysed in Triton lysis buffer (50 mM Tris pH 7.5, 1% Triton X-100, 150 mM NaCl, 50 mM NaF, 2 mM Na-vanadate, 0.011 g/ml beta-glycerophosphate), supplemented with 1x PhosSTOP phosphatase inhibitors (#04906837001, Roche) and 1x cOmplete protease inhibitors (#11697498001, Roche). For all experiments in which ubiquitination of mTOR was assessed, the lysis buffer was supplemented with 10 mM NEM (N-ethylmaleimide; #E3876, Sigma) as a DUB inhibitor. For the immunoprecipitation of mTOR, 1 µl of antibody was added to lysates and samples were incubated rotating for three hours at 4 °C. After that, 30 µl of protein A agarose beads (#11134515001, Roche), pre-equilibrated with IP wash buffer (50 mM Tris pH 7.5, 1% Triton X-100 or 0,3% CHAPS, 50 mM NaF, 150 mM NaCl), were added to the samples, which were incubated with rotation for an extra hour at 4 °C, followed by four washes with IP wash buffer. For the immunoprecipitation of FLAG-tagged proteins, lysates were incubated with 20 μl slurry of anti-FLAG M2 affinity gel (Sigma, #A2220), pre-equilibrated with IP wash buffer, rotating for two hours at 4 °C. Samples were then washed four times with IP wash buffer, and beads were boiled in 2x SDS sample buffer (6 min, 95 °C). Samples were analyzed by immunoblotting as indicated in the figures.

### Streptavidin pulldown

Cells transfected with the pcDNA4/TO/SBP-mTOR construct were lysed 48 hours post-transfection in Triton lysis buffer. After lysis, samples were incubated for two hours at 4 °C with 30 µl slurry of streptavidin-sepharose beads (#GE17-5113-01, Sigma), pre-equilibrated with IP wash buffer. Samples were then washed four times with IP wash buffer and beads were boiled in 2x SDS sample buffer (6 min, 95 °C).

### His-tag pulldown

For the pulldown of His_6_-tagged mTOR, cells were lysed in 300 µl binding buffer [1x PBS, 8 M urea (#15604, Sigma), 10 mM imidazole (#A1073, Applichem), 300 mM NaCl]. Lysates were sonicated four times for 15 seconds with 15-second breaks. The samples were then subjected to three freeze-thaw cycles (from liquid N_2_ to 37 °C), followed by centrifugation (17000 g, 5 min, RT) to clarify the lysates. Fifty µl of lysates were kept as input, and 12.5 µl of 6x SDS sample buffer were added before boiling. The remaining sample was incubated with 100 µl of Ni-NTA (nickel-nitriloacetic acid; #1018244, Qiagen) slurry, pre-equilibrated with binding buffer, for two hours at 4 °C. Beads were then washed five times with binding buffer, followed by two washes with binding buffer containing 30 mM imidazole, and boiling in 2x SDS sample buffer (6 min, 95 °C).

### *In vitro* deubiquitination assay

Cells transfected with pcDNA3-FLAG-mTOR were lysed 48 hours post-transfection with Triton lysis buffer without NEM. mTOR immunoprecipitation was performed as described above, with an additional wash with 50 mM Tris pH 7.5 after the washes with IP wash buffer. Subsequently, 200 ng of recombinant CYLD (#E-556, Boston Biochem) in 10 µl DUB reaction buffer (50 mM Tris pH 7.5, 5 mM DTT) were added to the beads. Samples were incubated at 37 °C for one hour and the reaction was terminated by addition of 2x SDS sample buffer and boiling (6 min, 95 °C).

### Proximity biotin labelling and streptavidin pulldown of interactors

To also capture weak or transient protein-protein interactions between mTOR and CYLD, in addition to more stable interactions, we utilized a H_2_O_2_-inducible, APEX2 proximity-based biotinylation system, as described in ^68^. In brief, cells expressing FLAG-APEX2-mTOR were labelled by the addition of 500 μM biotin-phenol (#LS-3500, Iris Biotech) to the medium for 30 min, followed by addition of 1 mM H_2_O_2_ for one min to induce the biotinylation reaction. Cells were then washed three times with quenching buffer [1x PBS, 10 mM sodium azide, 10 mM sodium ascorbate, 1 mM Trolox (6- hydroxy-2,5,7,8-tetramethylchroman-2-carboxylic acid) (#238813, Sigma-Aldrich)]. Cells were lysed with a modified RIPA buffer (50 mM Tris, 150 mM NaCl, 1% Triton X- 100, 0,5% sodium deoxycholate, 0.1% SDS in PBS), supplemented with 10 mM sodium ascorbate, 1 mM sodium azide, 1 mM Trolox, and 1x cOmplete protease inhibitors. Next, samples were incubated on ice for 15 min and centrifuged at 10000 g for 10 min. Protein concentration was measured using the Pierce 660 nm assay (#22660, Thermo Scientific) and equal protein amounts were used for the pulldowns. A portion of the lysate was kept aside as input and the remaining material was incubated with streptavidin-sepharose beads, pre-equilibrated with the modified RIPA lysis buffer, rotating for one hour at RT. The beads were washed five times with modified RIPA buffer and boiled in 2x SDS sample buffer (6 min, 95 °C).

### SUrface SEnsing of Translation (SUnSET) assays

To measure changes in *de novo* protein synthesis in cells upon CYLD knockdown or knockout, a puromycin incorporation assay was utilized. Cells were treated with Torin1 for 16 hours, cycloheximide (CHX) for four hours, or DMSO as control. Thirty minutes before lysis, puromycin (10 µg/ml) was added to the culture media. Cells were lysed as described above and lysates were subjected to SDS-PAGE for detection of puromycin incorporation into nascent polypeptide chains using an anti-puromycin antibody (Table S2). Signals were quantified using the GelAnalyzer 19.1 software. Protein synthesis levels were determined as the ratio of total puromycin signal to GAPDH intensity.

### Luciferase reporter assays

Changes in NF-κB pathway activation were assessed using a standard luciferase reporter assay. Cells were transfected with siRNAs targeting CYLD and IKKβ either alone or in combination as described above. On the next day, cells were transfected with 200 ng of the 3xKBL reporter construct (expressing firefly luciferase under the control of 3 tandem NF-κB binding sites) and 100 ng of the internal control pRL-null vector (expressing Renilla luciferase) using the Effectene reagent as described above. Forty-eight hours later, luciferase assays were performed with the Dual Luciferase Assay Reporter System (#E1910, Promega), as per the manufacturer’s instructions. NF-κB activation was calculated as the ratio of firefly luciferase to Renilla luciferase values.

### Cell size measurements

Changes in cell size were measured as a physiological readout downstream of mTORC1 activity in CYLD knockdown or knockout cells using an IncuCyte S3 live-cell imaging and analysis System (Sartorius). Transient knockdowns of TSC2 were used as a positive control in RNAi experiments, and Torin1 treatments were used to test the involvement of mTOR activation in cell size alterations. In brief, cells cultured on 12- well plates and images from nine different regions per well were acquired at regular intervals with a 10x objective in an IncuCyte apparatus. Images were analyzed using the IncuCyte software, and cell size was determined 66 hours after the initiation of the experiment.

### *C. elegans* strains and maintenance

*C. elegans* were grown and kept at 20 °C on standard Nematode Growth Medium (NGM) and fed *E. coli* (OP50) ^74^. Wild-type (N2) worms were obtained from the *Caenorhabditis* Genetics Center (CGC) (University of Minnesota), which is supported by the NIH Office of Research Infrastructure Programs (P40 OD010440). The VC533 *raga-1(ok701)* strain was acquired from the *C. elegans* Knockout Consortium (University of Oklahoma) via CGC. The AA4408 (*raga-1(ok701)*; sqIs17[*hlh-30p::hlh-30::*GFP + rol-6(su1006)]) strain was a kind gift by Adam Antebi (MPI-AGE).

### *C. elegans* lifespan experiments

The worm larvae were synchronized by egg laying protocol and grown onto plates with OP50 *E. coli* at 20 °C until they developed into L4 stage hermaphrodites. In all experiments, RNAi treatment was initiated after development. Once hermaphrodite worms reached to late L4 stage, they were transferred onto plates with HT115 *E. coli* carrying empty vector or *cyld-1* RNAi construct for lifespan assays. As an initial population, 96 worms were assessed per condition and scored every day or every other day at 20 °C ^75^. The worms lost or burrowed into the medium as well as those with ‘protruding vulva’ or that underwent bagging were censored.

### Subcellular localization of HLH-30::GFP analysis in *C. elegans*

The cyto-nuclear relocalization of GFP-tagged HLH-30 was used as a proxy for TORC1 activity in worms. To determine the localization of HLH-30::GFP, AA4408 (*raga-1(ok701)*; sqIs17[*hlh-30p::hlh-30::*GFP + rol-6(su1006)]) worms were used. The synchronized worms were transferred onto plates with HT115 *E. coli* carrying empty vector or *cyld-1* RNAi construct after they reached into L4 stage hermaphrodites. At day five of adulthood, the worms were immobilized using 0,1% azide in M9 buffer on 2% agarose pads to visualize nuclear localization of HLH-30::GFP. Fluorescence images were taken with an Axio Imager Z1 microscope (Zeiss).

### Human sample collection

Normal human skin samples and skin tumor samples were obtained from Department of Dermatology, University of Cologne (CCS patient #2 and control) and the Department of Medical Genetics, Faculty of Medicine, University of Szeged, Szeged, Hungary (CCS patient #1). The *CYLD* gene mutation in patient #1 (CYLD c.2797C/T, p.Arg933Ter) was identified by genomic DNA sequencing. Both patients were diagnosed positive for CCS based on standard diagnostic and histological criteria. Patient consent was obtained from all human subjects and all procedures were performed in accordance with the Declaration of Helsinki.

### Hematoxylin and eosin (H&E) staining and Immunohistochemistry

Skin biopsies were fixed in 4% paraformaldehyde (PFA), embedded in paraffin, and 10 µm sections were prepared. For H&E staining, paraffin was removed and staining was performed using the automated Gemini slide stainer (Thermo). Stained tissues were mounted on No. 1 rectangular coverslips. Images of H&E-stained tissue samples were obtained using a 10x objective on a Zeiss Axioscan 7 microscope.

For immunohistochemistry staining, paraffin sections were deparaffinized and incubated with citrate buffer (pH 6.0) for one hour at 98 °C to retrieve antigen. Sections were further incubated with 3% hydrogen peroxide for 10 min to block endogenous peroxidase activity and 10% goat serum in PBS to block unspecific binding sites. Next, the sections were incubated with anti-phospho-S6 antibody (Ser240/244) (#5364, Cell Signaling Technology) or anti-phospho-4E-BP1 (Thr37/46) (#2855, Cell Signaling Technology) overnight at 4 °C. Subsequently, the sections were incubated with HRP- conjugated anti-rabbit antibody (#K4003, Dako) for one hour at RT. The slides were washed 3x with 1x PBS after antibody incubations. Signals were detected with 3,3’- diaminobenzidine substrate and the sections were counterstained with hematoxylin and mounted. Images were obtained using a 10x objective on a Zeiss Axioscan 7 microscope.

## Statistical analysis

Statistical analysis and presentation of quantification data was performed using GraphPad Prism (versions 9.1.0 and 9.2.0). Data in graphs are shown as mean ± SEM. The normality of data distribution was tested using the Shapiro–Wilk and Kolmogorov– Smirnov tests in Prism. For graphs with only two conditions shown and normal data distribution, significance for pairwise comparisons was calculated using Student’s *t*- tests. For graphs with only two conditions shown and non-normal data distribution, significance for pairwise comparisons was calculated using Mann–Whitney *U* tests. For graphs with three or more conditions shown, significance for pairwise comparisons to the respective controls was calculated using one-way analysis of variance (ANOVA) with post hoc Holm–Sidak test. Sample sizes (n) and significance values are indicated in the figure legends (* p < 0.05, ** p < 0.01, *** p < 0.001, **** p < 0.0001, ns: non-significant).

For worm lifespan experiments, median lifespan was determined with GraphPad Prism 6.0 software which was also used to generate lifespan graphs. The mean lifespan was determined with OASIS software ^76^. Significance was calculated using the log-rank (Mantel-Cox) method for the comparison of two survival functions through the complete lifespan experiment. For the analysis of HLH-30::GFP localization in *C. elegans*, significance was assessed using two-way ANOVA.

All findings were reproducible over multiple independent experiments, within a reasonable degree of variability between replicates. The number of replicate experiments for each assay is provided in the respective figure legends. No statistical method was used to predetermine sample size, which was determined in accordance with standard practices in the field. No data were excluded from the analyses. The experiments were not randomized, and the investigators were not blinded to allocation during experiments and outcome assessment.

## Acknowledgements

We thank all members of the Demetriades lab for critical discussions; Andreas Lamprakis for technical support with cloning; and the MPI-AGE FACS & Imaging Core Facility for support with microscopy. SAF and JP received support by the Cologne Graduate School of Ageing Research. CD is funded by the European Research Council (ERC) under the European Union’s Horizon 2020 research and innovation programme (grant agreement No 757729), and by the Max Planck Society. Parts of this work were supported by the ERC under the European Union’s Seventh Framework Programme via an ERC Starting Grant (grant agreement No 260602) to AAT; and by the Deutsche Forschungsgemeinschaft (DFG, German Research Foundation) through the Research Unit Grant FOR2722 (DE 3170/1-1; Project No 384170921) to CD, and the DFG grant VI 742/4-2 to DV. Graphical models in figures were created with BioRender.com.

## Author Contributions

Experimental work: SAF, JP, DST, SW, JN; data analysis: SAF, CD; worm experiments and data analysis: SK, DV; human material collection, processing and analysis: SAF, XD, IBN, SA-G, MS, SAE; resources: CG, GM, MP; project design: SAF, CD; conceptualization: SAF, GM, MP, AAT, DV, CD; project supervision: CD; funding acquisition: AAT, SE, DV, CD; figure preparation: SAF, CD; manuscript draft: SAF, CD, with contributions from all authors. All authors approved the final version of the manuscript and agree on the content and conclusions.

## Declaration of interests

The authors declare no competing interests.

## Data availability

The data that support the findings of this study (uncropped immunoblots, microscopy pictures) are available from the corresponding authors upon reasonable request. All unique plasmids and cell lines generated in this study are available from the corresponding authors on reasonable request, with a completed material transfer agreement.

## Code availability

No code was generated in this study.

## Additional Information

Supplemental Information (Figures S1-6 and Tables S1-3) is available for this paper.

Correspondence and requests for materials should be addressed to Constantinos Demetriades or Stephanie A. Fernandes.

## Notes

### Competing Interest Statement

The authors have declared no competing interest.

