## Supplementary material for "The tumor suppressor CYLD acts as a deubiquitinase for mTOR to constrain its activity": Figures S1-S6

### Supplemental Information

Tables S1-3 and Figures S1-6

### Supplemental Tables

Table S1. RNAi screen for DUB genes as putative mTORC1 regulators.

Table S2. List of primary antibodies used in this study.

Table S3. List of DNA oligonucleotides and donor templates used in this study.

### Supplemental Figures

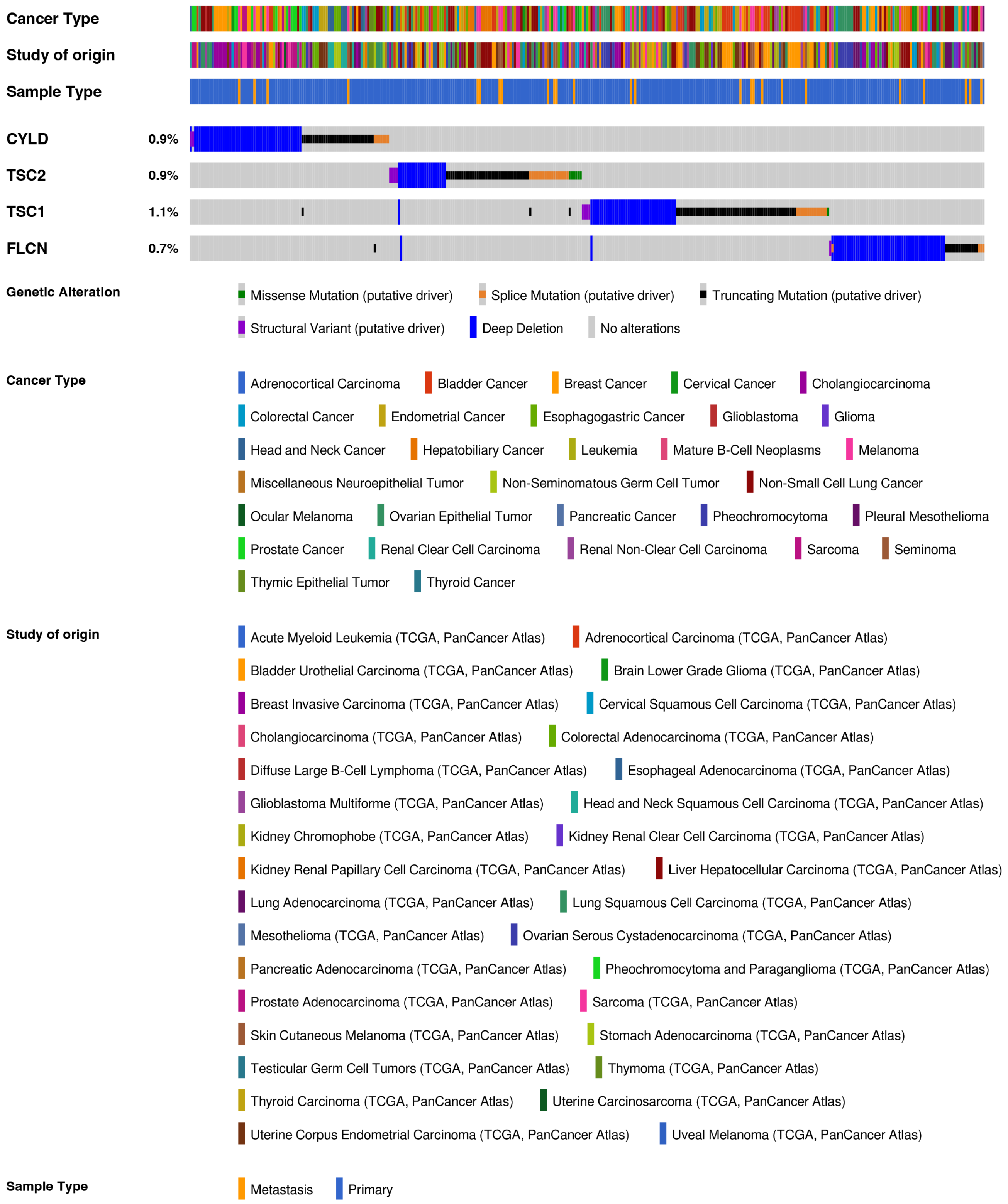


**Figure S1. *CYLD* mutations do not coincide with mutations in other negative mTORC1 regulator genes.**

Analysis of mutations and copy number alterations in the *CYLD*, *TSC2*, *TSC1* and *FLCN* genes analyzed in diverse cancer types using the cBioPortal platform for Cancer Genomics (www.cbioportal.org) shows a largely mutual exclusive relationship between *CYLD* and the other interrogated genes, suggesting it may be acting as a negative upstream regulator in the mTORC1 signaling pathway. See Methods for additional information.

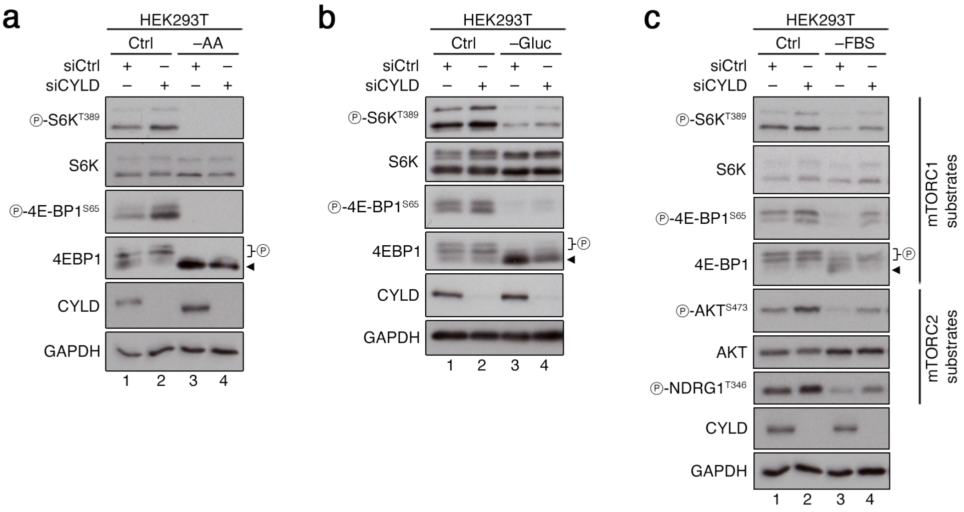


**Figure S2. mTORC1 activity is responsive to nutrient or growth factor removal in CYLD knockdown cells.**

**(a)** Immunoblots with lysates from HEK293T cells transiently transfected with siRNAs targeting *CYLD* or a control RNAi duplex (siCtrl), treated with media containing or lacking AAs (–AA) for 1 h before lysis, probed with the indicated antibodies. n = 3 independent experiments.

**(b)** As in (a), but for glucose starvation (–Gluc) for 1 h before lysis. n = 3 independent experiments.

**(c)** As in (a), but for growth factor starvation (–FBS) for 16 h before lysis. n = 3 independent experiments.

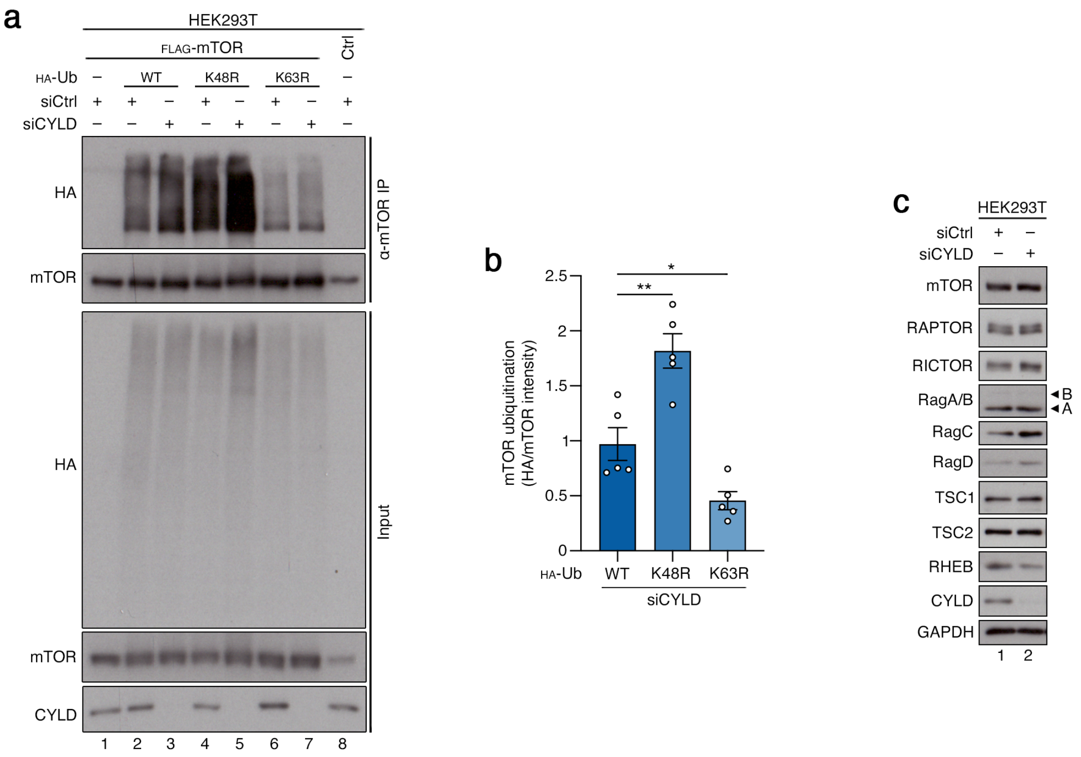


**Figure S3. CYLD removes non-degradative, K63-linked ubiquitin chains from mTOR, without consistently affecting the protein levels of most mTOR pathway components.**

**(a-b)** Analysis of mTOR ubiquitination in HEK293T cells transiently expressing FLAG-tagged mTOR and HA-tagged WT, K48R or K63R Ub, with or without siRNA-mediated knockdown of *CYLD*. Ubiquitination of immunopurified mTOR was assessed by immunoblotting with the indicated antibodies (a). Quantification of mTOR ubiquitination with WT or mutant Ub molecules in the siCYLD samples shown in (b). Data shown as mean ± SEM. * p<0.05, ** p<0.01. n = 5 independent experiments.

**(c)** *CYLD* knockdown does not affect the protein levels of most mTOR pathway components. Immunoblots with lysates from HEK293T cells, transiently transfected with siRNAs targeting *CYLD* or a control RNAi duplex (siCtrl), probed with the indicated antibodies. n = 3 independent experiments.

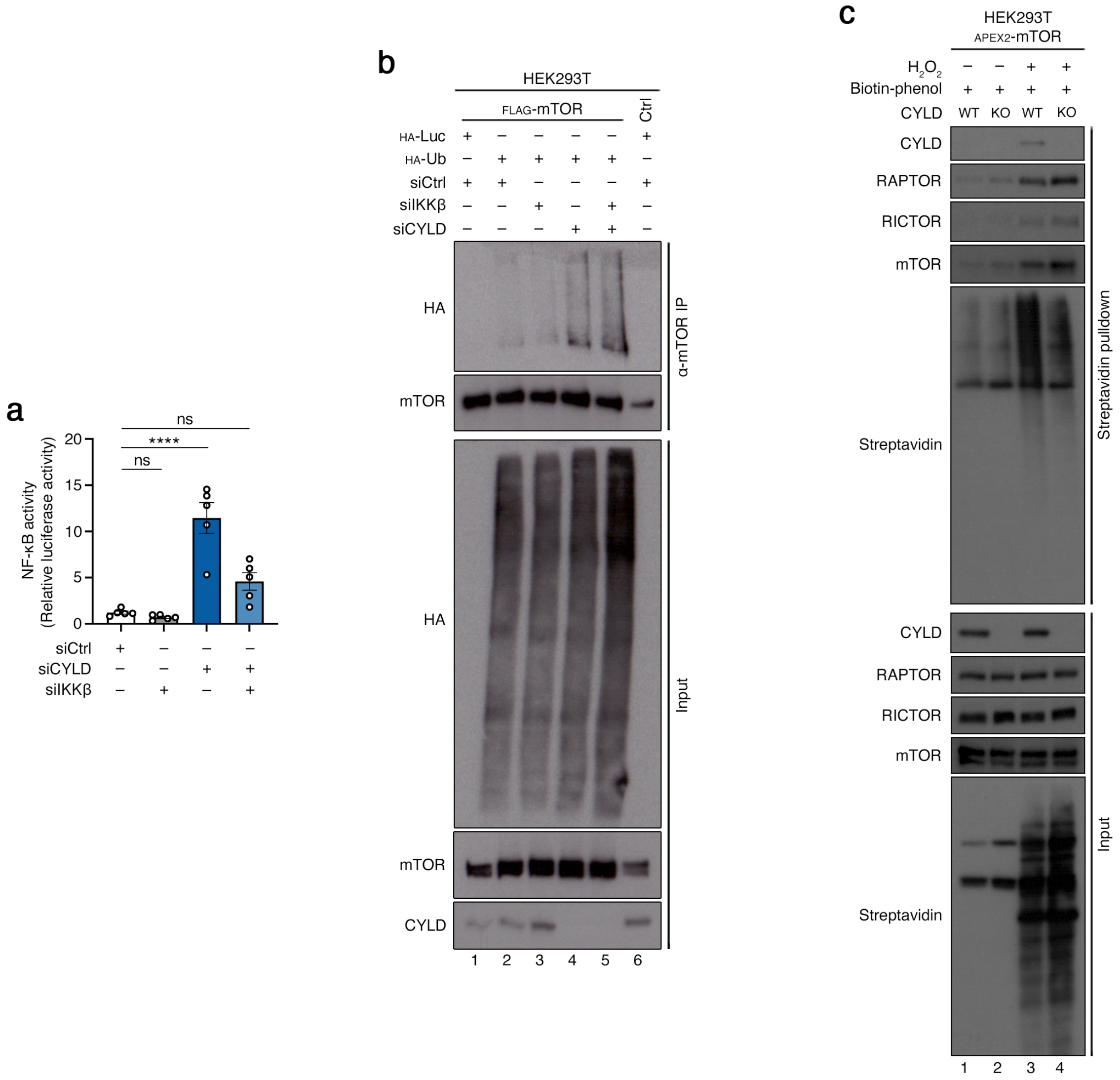


**Figure S4. CYLD interacts with mTOR and controls its ubiquitination independently of its canonical role in NF-κB signaling.**

**(a)** A transcriptional NF-κB reporter assay confirms the positive and negative roles of IKKβ and CYLD, respectively, in NF-κB signaling. Dual luciferase reporter assays for NF-κB activation using lysates from HEK293T cells transfected with siRNAs targeting *CYLD*, *IKKβ* or a control RNAi duplex (siCtrl). n = 5 independent experiments. Data shown as mean ± SEM. **** p<0.0001, ns: non-significant.

**(b)** CYLD regulates the ubiquitination of mTOR independently of changes in NF-κB signaling activity. Analysis of mTOR ubiquitination in HEK293T cells transiently expressing FLAG-tagged mTOR and HA-tagged Ub, transfected with siRNAs targeting *CYLD*, *IKKβ* or a control RNAi duplex (siCtrl). Ubiquitination of immunopurified mTOR was assessed by immunoblotting with the indicated antibodies. n = 3 independent experiments.

**(c)** Proximity biotinylation assay identifies CYLD as an mTOR interactor. Immunoblots with lysates and streptavidin pulldowns from HEK293T cells stably expressing FLAG-APEX2-

mTOR probed with the indicated antibodies. The respective CYLD KO HEK293T cells were used as a control for signal specificity. Protein biotinylation in the vicinity of mTOR was induced by acute addition of $H_2O_2$ before lysis. n = 3 independent experiments.

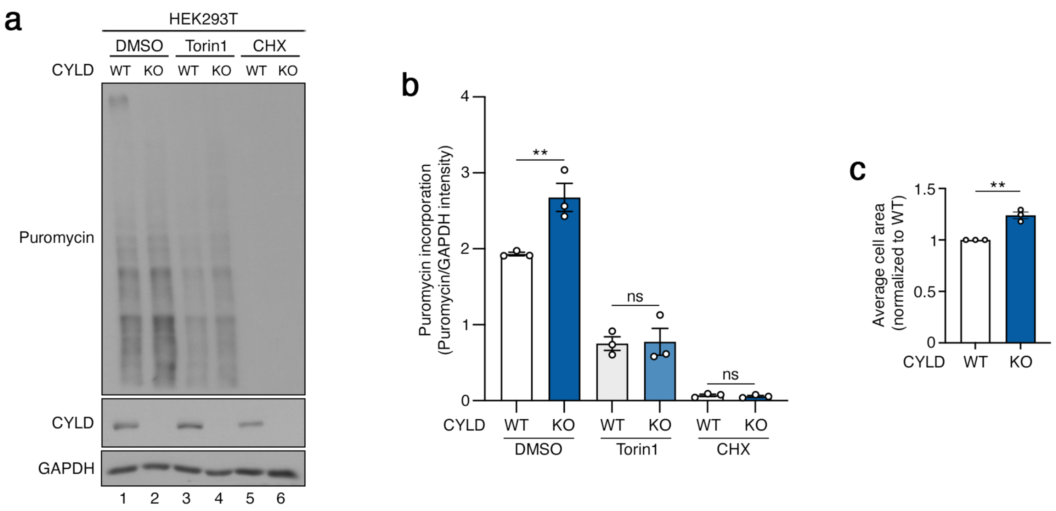


**Figure S5. CYLD loss leads to increased protein synthesis and cell size.**

**(a-b)** Elevated translation rates in CYLD KO cells are reversed by mTOR inhibition. Puromycin incorporation assays in WT or CYLD KO HEK293T cells treated with Torin1 (250 nM, 16 h), CHX (100 µM, 4 h), or DMSO as control, analyzed by immunoblotting with the indicated antibodies (a). Quantification of puromycin incorporation into nascent polypeptide chains in (b). n = 3 independent experiments.

**(c)** Increased cell size of CYLD knockout cells is reversed. Cell area measurements from WT or CYLD KO HEK293T cells using an IncuCyte S3 live-cell imaging and analysis system. n = 3 independent experiments.

Data in graphs shown as mean ± SEM. ** $p<0.01$, ns: non-significant.

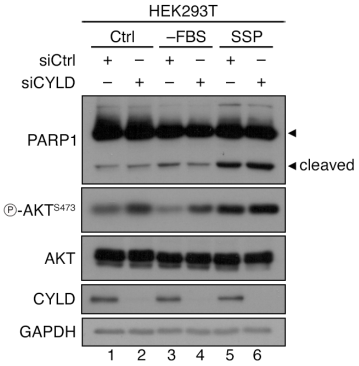


**Figure S6. CYLD knockdown blunts the serum-starvation-induced cell death downstream of mTORC2 inactivation.**

Immunoblots with lysates from HEK293T cells transiently transfected with siRNAs targeting *CYLD* or a control RNAi duplex (siCtrl), treated with media containing or lacking growth factors (–FBS) for 24 h before lysis, probed with the indicated antibodies. PARP1 cleavage was used as a proxy for cell death induction. Staurosporine (SSP; 1 μM, 6 h) was used a positive control. n = 3 independent experiments.
